# Dose-dependent functions of SWI/SNF BAF in permitting and inhibiting cell proliferation *in vivo*

**DOI:** 10.1101/636720

**Authors:** Aniek van der Vaart, Molly Godfrey, Vincent Portegijs, Sander van den Heuvel

## Abstract

SWI/SNF complexes regulate transcription through chromatin remodeling and opposing gene silencing by Polycomb-group (PcG) proteins. Genes that encode SWI/SNF subunits are frequently mutated in human cancer. The selective advantage, subunit bias, and common heterozygosity of such mutations remains poorly understood. Here, we characterized how functional loss of various SWI/SNF subunits and PcG EZH2 affect proliferation-differentiation decisions *in vivo*, making use of the reproducible development of the nematode *C. elegans.* We applied a lineage-specific genetics strategy to create partial or complete SWI/SNF subunit loss, as well as double gene knockout with PcG EZH2. Our data show that a high SWI/SNF BAF dosage is needed to oppose Polycomb-mediated transcriptional repression and to arrest cell division during differentiation. In contrast, even in the absence of the PcG EZH2-related methyltransferase, a low level of the SWI/SNF BAF complex is necessary and sufficient to sustain cell proliferation and hyperplasia. Our data provide experimental support for the theory that during carcinogenesis partial SWI/SNF BAF loss-of-function mutations are selected because they eliminate a tumor suppressor activity while maintaining an essential transcription regulatory function.

## Introduction

During development and tissue homeostasis, stem and progenitor cells proliferate to generate daughter cells that acquire specialized functions. The terminal differentiation of such cells coincides with a permanent withdrawal from the cell division cycle. This cell-cycle arrest is achieved by a combination of cell cycle regulators that include the retinoblastoma tumor suppressor (Rb) protein family of transcriptional co-repressors, CDK-inhibitory proteins (CKIs) that bind and block cyclin-dependent kinases (CDKs), and the anaphase-promoting complex in association with the coactivator Cdh1/FZR1 (APC/C-FZR1), which promotes protein degradation (Ruijtenberg and van den Heuvel 2016). In addition to these general regulators of the cell cycle, lineage-specific transcription factors and chromatin regulators coordinate the arrest of cell division with terminal differentiation. In particular, SWI/SNF (switch/sucrose non-fermenting) chromatin remodeling complexes have been found to play an important role in this process (Ruijtenberg and van den Heuvel 2015; Yu et al. 2013; Joliot et al. 2014; Albini et al. 2015).

SWI/SNF chromatin remodeling complexes are large, multi-subunit protein complexes, initially identified as positive regulators of gene expression in yeast (for review, (Mathur and Roberts 2018)). The conserved components of SWI/SNF complexes were independently identified in *Drosophila melanogaster* as antagonists of Polycomb-mediated transcriptional repression, and found in mammals through homology searches. Multiple distinct SWI/SNF subcomplexes are modularly assembled from a variety of different subunits. These SWI/SNF complexes contain an ATPase core subunit and use the energy generated by ATP hydrolysis to alter nucleosome occupancy at gene regulatory regions, and to evict Polycomb-repressor complexes (Kadoch and Crabtree 2015; Wilson and Roberts 2011; Kadoch et al. 2016). SWI/SNF complexes can be divided into BAF and PBAF variants, as well as a recently identified non-canonical ncBAF complex (Mashtalir et al. 2018). Canonical SWI/SNF complexes consist of one of two mutually exclusive ATPase subunits (BRM/SMARCA2 or BRG1/SMARCA4), highly conserved ‘core’ subunits (SNF5/SMARCB1, BAF155/SMARCC1, BAF170/SMARCC2), an array of variable ‘accessory’ subunits, and BAF or PBAF-specific variant subunits (respectively ARID1A/B, and ARID2, PBRM1, BRD7) (Mashtalir et al. 2018) (figure supplement 1a-b). How the divergence in SWI/SNF subcomplexes and variant subunits relate to function is not clear, although this variation most likely plays a role in the regulation of lineage-specific functions of the complexes (Kadoch and Crabtree 2015; Wilson and Roberts 2011; Kadoch et al. 2016).

Importantly, mammalian SWI/SNF complexes act as tumor suppressors and are altered in a wide variety of cancers. In fact, mutations in the collective set of SWI/SNF subunit-encoding genes have been found in 20% of the examined human cancers (Wilson and Roberts 2011; Shain and Pollack 2013; Kadoch et al. 2013). The broad spectrum of the identified genetic alterations makes it difficult to understand the exact oncogenic effects. While the BAF-specific subunit ARID1A is most frequently mutated, alteration of specific SWI/SNF subunits is associated with specific cancer types (Wilson and Roberts 2011; Shain and Pollack 2013; Kadoch et al. 2013). Cancer-associated SWI/SNF missense mutations or deletions are often heterozygous, or affect subunits for which paralogues exist (Kadoch and Crabtree 2015; Bögershausen and Wollnik 2018). This can lead to tumor dependence on the paralogous protein, as revealed by requirement for ARID1B in ARID1A mutant cancers, or BRM (SMARCA2) in tumors with mutations in the BRG1 (SMARCA4) ATPase, in synthetic lethal screens (Helming et al. 2014b; Hoffman et al. 2014; Wilson et al. 2014). Heterozygous mutations in genes encoding SWI/SNF subunits are also associated with intellectual disability disorders such as Coffin-Siris syndrome (Bögershausen and Wollnik 2018). The prevalence of genetically dominant SWI/SNF gene mutations in cancer and neurologic diseases indicate dosage dependent functions of SWI/SNF complexes.

Several mechanisms have been proposed to explain the remarkable spectrum of SWI/SNF gene mutations. The correlation between cancer type and mutated subunits has been taken to indicate that specific subunits protect against cancer in specific tissues (Kadoch et al. 2013). Mutation or deletion of specific SWI/SNF subunits could lead to the formation of complexes with alternative composition, thereby removing a tumor suppressive function and activating a pro-oncogenic transcription program by the remaining, perhaps aberrantly assembled, SWI/SNF complexes (Helming et al. 2014b; Wang et al. 2009, 5). Alternatively, various subunits of the SWI/SNF complex may be haploinsufficient, implying a dosage-dependent tumor suppressor role for the complex (Kadoch and Crabtree 2015; Wilson and Roberts 2011; Kadoch et al. 2016). In this model, a reduction of the tumor suppressive function of SWI/SNF would sufficiently alter gene expression patterns to predispose cells to tumor development (Wilson and Roberts 2011; Kadoch et al. 2016). In certain cases, mutations appear to lead to neomorphic gain-of-function or dominant-negative inhibition of SWI/SNF complexes and thereby promote cancer formation or neurologic disease. For instance, point mutations in the ATPase domain can lead to assembly of non-functional complexes, which changes chromatin accessibility by competing with wild type complexes (Bögershausen and Wollnik 2018; Hodges et al. 2018). Whether the observed SWI/SNF subunit mutations reflect neomorphic or dominant-negative effects, haploinsufficient functions, or cancer-type specific variations of these models, remains a topic of investigation, as do the reasons behind the strong cancer-type bias in mutated subunits.

In this study, we characterize how partial versus complete loss-of-function of various SWI/SNF subunits affect the *in vivo* proliferation and differentiation of muscle precursor cells. We take advantage of the invariant cell lineage and advanced possibilities for controlled manipulation in the nematode *Caenorhabditis elegans* (Sulston and Horvitz 1977). Using lineage-specific gene knockout and protein degradation technologies, we demonstrate that core subunits of the SWI/SNF BAF complex contribute strong dosage-dependent functions in cell proliferation. As such, partial loss of function of BAF subunits leads to hyperplasia, which is enhanced by loss of negative cell-cycle regulators. This indicates a tumor suppressive function of SWI/SNF BAF, which resides in part on PcG protein opposition. Strikingly, we found that in the same cells, low levels of the SWI/SNF complex are required for cell proliferation, independently of the presence of PcG proteins or negative cell-cycle regulators. Our single molecule FISH and RNA-sequencing studies show that acute inactivation of SWI/SNF BAF in muscle precursor cells rapidly alters the transcript levels of several hundred genes, including cyclin D, demonstrating that the complex is continuously required for the regulation of gene expression. Thus, in the same cell type and developmental decisions, a high dosage of SWI/SNF BAF subunits is needed for temporal arrest of cell division and PcG opposition, while a low level is required to sustain proliferation. We propose that similar dosage-dependent effects explain the selection of SWI/SNF partial loss-of-function mutations during carcinogenesis.

## Results

### The SWI/SNF BAF subcomplex is crucial for cell division arrest during development

To investigate how the SWI/SNF complex regulates cell proliferation, we exploited the fact that cell divisions in the nematode *Caenorhabditis elegans* follow a well-characterized invariant pattern throughout development. Abnormal cell division patterns resulting from aberrant regulation of proliferation-differentiation processes can therefore be readily recognized, monitored and quantified based on *in vivo* observations. Previously, we observed that a lineage-specific temperature-sensitive mutation in the SWI/SNF core subunit gene *swsn-1* (SMARCC1/2) gives rise to hyperplasia during *C. elegans* post-embryonic mesoderm development. Importantly, this mutation combined with loss of negative cell-cycle regulators induces a unique tumorous overproliferation phenotype (Ruijtenberg and van den Heuvel 2015).

To examine the role of specific SWI/SNF subunits in the regulation of proliferation, we performed RNA interference (RNAi) experiments for *C. elegans* genes predicted to encode components of two distinct subcomplexes, BAF and PBAF. These complexes share core subunits and several additional proteins, while differing in specific factors (figure supplement 1a-b). We focused on the mesoblast (M) lineage, which includes two sequential periods of cell-cycle quiescence, proliferation and muscle differentiation (Sulston and Horvitz 1977) (Figure 1a). Integration of a “tagBFP2-to-mCherry” lineage-tracing reporter and an *hlh-8* Twist promoter-CRE recombinase transgene (P*hlh-8::CRE*) facilitated quantification of mCherry-positive mesoblast daughter cells (Figure 1b-c). Using this background, we observed that knockdown of the core ATPase subunit *swsn-4* BRM/BRG1, the core subunit *snfc-5* SNF5, as well as the BAF-specific subunit *swsn-8* ARID1 increased the number of M descendants. Knockdown of three PBAF specific SWI/SNF subunits only had weak effects (Figure 1d-f and figure supplement c-d). Simultaneous inhibition of negative-regulators of the cell cycle further emphasized the different contributions of BAF versus PBAF subunits. Single knockout of the APC/C activator *fzr-1* Cdh1, an inhibitor of cell-cycle entry, did not alter the M lineage division pattern. However, *fzr-1* loss enhanced the hyperplasia of M descendants when combined with knockdown of SWI/SNF core subunits and *swsn-8* ARID1, but not when combined with PBAF-specific subunits (Figure 1d-e and figure supplement 1c-d). These data indicate that the SWI/SNF BAF complex contributes critically to the cell division arrest of muscle precursor cells.

**Figure 1.**
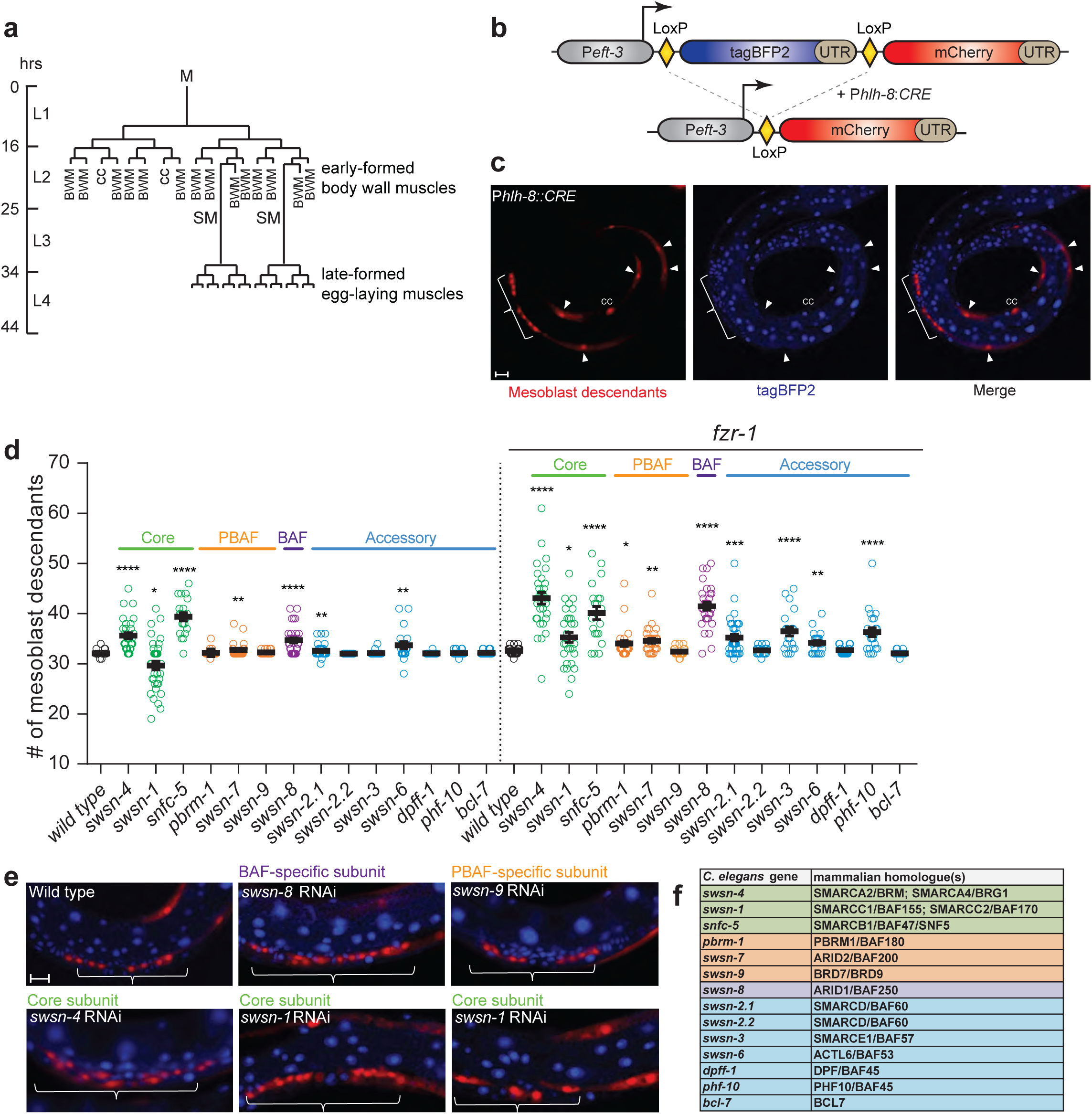
The SWI/SNF BAF complex promotes cell cycle exit. (**a**) Lineage of the *C. elegans* mesoblast (M), the single precursor cell of post-embryonic mesoderm. The M cell is born during early embryogenesis and initiates proliferation half way through the first larval stage (L1), to form 14 striated muscle cells (body wall muscle, BWM), two scavenger cells (coelomocytes, CC), and two new ventral muscle precursor cells (sex myoblasts, SM). The SMs remain quiescent while they migrate anteriorly to align with the vulva, resume proliferation late in the third larval stage (L3), and differentiate to form 16 muscle cells that are required for egg laying. (**b**) Design of the lineage-tracing reporter, which is single-copy integrated into the *C. elegans* genome. A universal promoter (P*eft-3*) drives expression of tagBFP2 flanked by two LoxP sites and followed by the *lin-858* UTR. Excision of tagBFP2 leads to mCherry expression, which provides an easily visible switch from blue-to-red fluorescence in cells where CRE is expressed and all daughter cells. (**c**) Representative image of mesoblast lineage descendants as marked by the lineage tracing construct in an L4 larva (lateral view, ventral down; arrowheads point to BWM, brackets indicate late egg-laying muscle precursors). (**d**) Quantification of total number of mesoblast lineage descendants per animal at the L4 larval stage following RNAi by feeding of synchronised L1 larvae for the indicated genes in wild type or *fzr-1* mutant backgrounds. (**e**) Representative images of the vulva region of RNAi treated larvae. Anterior to the left, ventral down, scale bar 10 µm in all images. (**f**) The SWI/SNF complex consists of core subunits sufficienct for *in vivo* chromatin remodelling (green), accessory (blue), and BAF (purple) and PBAF-specific (orange) signature subunits. Table of *C. elegans* names used in this publication and commonly used mammalian homologue names for the different subunits

### SWI/SNF gene knockout leads to two divergent cell proliferation phenotypes

RNAi of the third core subunit, *swsn-1* (SMARCC1/2), led to a surprisingly variable number of mesoblast descendants, with animals showing a range from fewer to more than the normal number of cells (Figure 1d-e and figure supplement 1c-d). To test whether this reflects variability in RNAi-induced loss of function, we created conditional knockout alleles, as SWI/SNF null mutations are lethal. Using CRISPR/Cas9-mediated genome editing, we introduced Lox sites in endogenous genes encoding the SWSN-1 and SWSN-4 core components, the BAF-specific SWSN-8 subunit and the accessory subunit SWSN-2.1 BAF60 (Figure 2a). We combined these loxed SWI/SNF alleles with the P*hlh-8::CRE* and tagBFP2-to-mCherry integrated transgenes, to induce M lineage specific gene deletion and reporter expression.

**Figure 2.**
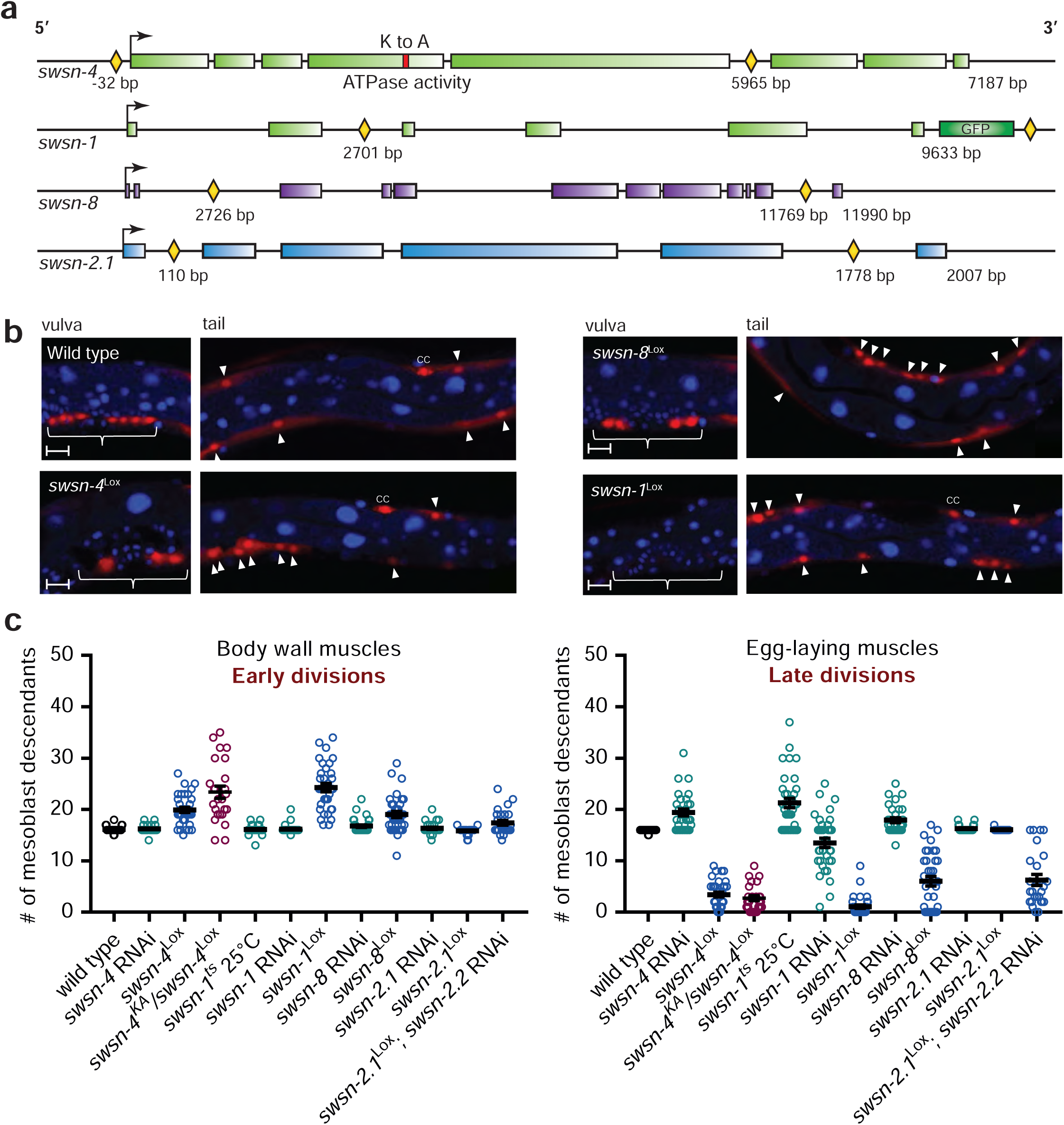
Knockout of endogenous SWI/SNF genes and RNAi induce highly divergent cell division phenotypes. (**a**) Schematic of Lox sites (yellow diamonds) integrated into the endogenous SWI/SNF genes indicated. The *swsn-4* ATPase-dead lysine to alanine mutation (K to A) is shown as a red block. Transcriptional start sites are indicated with arrows, introns are shown as black lines and exons as coloured blocks. (**b**) Representative images of mesoblast lineage descendants in wild type and indicated SWI/SNF gene knockout animals. Arrowheads point at BWM, brackets indicate egg-laying muscle precursors. Scale bar 10 µm. (**c**) Quantifications of mesoblast lineage descendants for the indicated genotypes, in the tail area (early dividing body wall muscles) and around the vulva (late dividing egg-laying muscles). Note that, in contrast to RNAi, SWI/SNF gene knockouts lead to overproliferation of the early dividing body wall muscle precursors, and cell division arrest of the late dividing egg-laying muscle precursor cells.

Remarkably, the *swsn-1, swsn-4*, and *swsn-8* knockout phenotypes differed greatly from those resulting from RNAi knockdown of the same genes, and from the temperature-sensitive *swsn-1* phenotype (Ruijtenberg and van den Heuvel 2015) (Figure 2b-c and figure supplement 2). Specifically, instead of the RNAi-induced extra M descendants in the vulva region, the gene knockouts resulted in fewer late muscle precursors. Moreover, early-formed M cell descendants, which exited the cell cycle normally upon RNAi treatment, overproliferated in the knockout strains (Figure 2b-c). Simultaneous inactivation of the *fzr-1* cell-cycle inhibitor synergistically increased the number of extra M lineage divisions in early development, but did not suppress the reduced number of late M lineage divisions (figure supplement 2). Thus the knockout data indicate that the SWI/SNF BAF complex exerts a critical function needed for cell number expansion, in addition to promoting cell-cycle arrest and differentiation.

In contrast to the other conditional SWI/SNF mutants, *swsn-2.1* knockout larvae remained normal. Two paralogous *C. elegans* genes, *swsn-2.1* and *swsn-2.2*, encode BAF60-related SWI/SNF subunits, compared to three paralogues in mammals (Ertl et al. 2016). When combined with *swsn-2.2* RNAi, the *swsn-2.1* knockout closely resembled the other conditional SWI/SNF gene knockouts (Figure 2c). This indicates that *swsn-2.1* and *swsn-2.2* BAF60 act redundantly, and likely in combination with core subunits as well as SWSN-8 ARID1, in proliferation control (Figure 2c).

### ATPase activity of the SWI/SNF complex is required both for cell division arrest and cell proliferation

To assess whether the knockout phenotypes result from loss of the ATPase-dependent functions of the complex, we created an ATPase-dead *swsn-4* allele by introducing a lysine-to-alanine (KA) mutation of a conserved residue that is essential for ATP hydrolysis (Richmond and Peterson 1996) (Figure 2a and figure supplement 2a). Because this mutant dies soon after hatching, we maintained the *swsn4*^KA^ mutation in a trans-heterozygous combination with a wild type or *swsn-4*^Lox^ allele. Following M lineage specific CRE expression, the *swsn-4*^KA^/*swsn-4*^Lox^ mutant showed similar or somewhat stronger cell-division abnormalities, compared to homozygous *swsn-4*^Lox^ knockout animals (Figure 2c and figure supplement 2). These data support that ATPase activity of the SWI/SNF BAF complex is required to promote both the cell-cycle arrest of early body wall muscle (BWM) precursors and the expansion of egg-laying muscle precursor cells in late development.

### SWI/SNF BAF subunit loss-induced hyperplasia coincides with delayed differentiation

We used promoter-fusion reporters and single-molecule FISH (smFISH) experiments to examine the proliferation-differentiation status of the SWI/SNF knockout cells. This showed residual *hlh-8* Twist expression, which is normally restricted to undifferentiated muscle precursors (Harfe et al. 1998) (figure supplement 3a-b). Moreover, expression of the differentiation-specific *myo-3* myosin heavy chain reporter was reduced at the time of normal BWM differentiation (Fire et al. 1998) (figure supplement 3c-d). Further, expression of the S-phase cyclin *cye-1* cyclin E was increased and expression of the CDK inhibitor *cki-1* Kip1 decreased compared to wild type, based on quantification of the number of mRNA copies/cell in smFISH experiments (figure supplement 3e). These data support that the extra cells in the conditional SWI/SNF knockout strains result from a prolonged proliferative undifferentiated state compared to wild type mesoblast descendants.

### SWI/SNF BAF-mediated cell-cycle arrest involves opposing Polycomb repression

SWI/SNF complexes oppose gene silencing by Polycomb repressor complexes PRC1 and PRC2 (Kennison and Tamkun 1988; Tamkun et al. 1992; Kia et al. 2008). We wondered whether unrestrained PcG-mediated gene silencing underlies the early overproliferation as well as the late cell-division arrest phenotypes of SWI/SNF knockout mesoblast daughters. Making use of GFP-tagged endogenous genes, we observed that SWI/SNF BAF subunits and the MES-2 EZH2 (PRC2) protein are expressed in the M lineage and throughout *C. elegans* development (figure supplement 4a-b). We combined mesoblast-specific knockout of *mes-2* EZH2 and various SWI/SNF complex subunits to determine whether the SWI/SNF knockout phenotypes depend on PcG-mediated gene silencing (Figure 3a-b). Similar to RNAi (Ruijtenberg and van den Heuvel 2015), the knockout of *mes-2* substantially reduced the overproliferation of early muscle precursor cells in SWI/SNF gene-knockout animals. The double knockout often showed close to wild type BWM numbers (Figure 3b). Contrary to this early effect, knockout of *mes-2*^Lox^ did not suppress the arrest of late mesoblast descendants in SWI/SNF mutants (Figure 3b). In fact, the removal of *mes-2* exacerbated the cell division arrest of SWI/SNF mutant late egg-laying muscle precursor cells (Figure 3b). Thus, to promote cell-cycle arrest and differentiation of early muscle precursors, SWI/SNF BAF apparently antagonizes Polycomb-mediated transcriptional repression, while the requirement for SWI/SNF BAF detected at later stages involves a mechanism other than PcG opposition.

**Figure 3.**
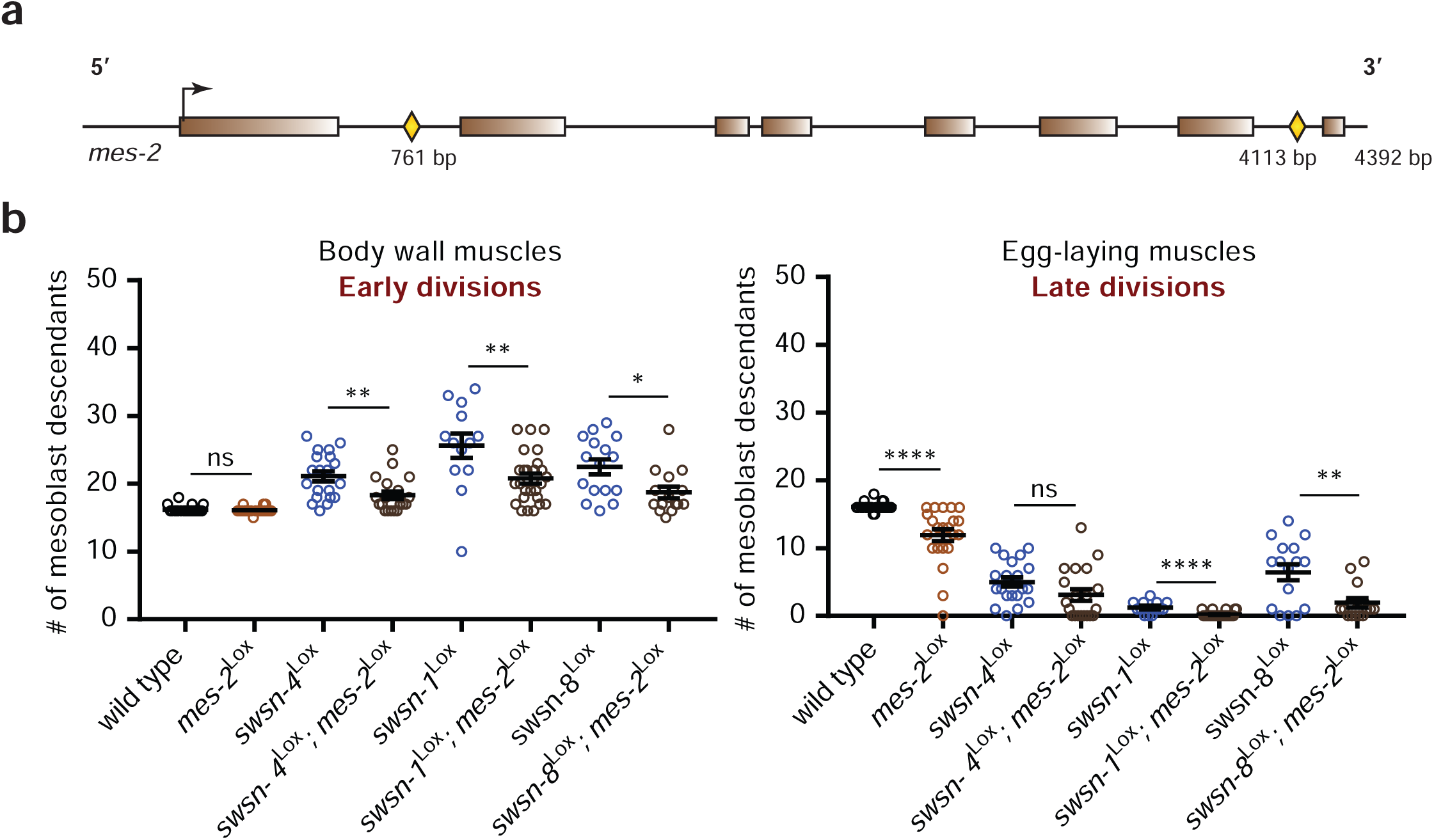
Simultaneous knockout of Polycomb PRC2 and SWI/SNF subunit genes rescues mesoblast descendant overproliferation, but not cell division arrest. (**a**) Schematic of Lox site integrations into the endogenous EZH2-related Polycomb gene *mes-2*, with Lox sites indicated by yellow diamonds. The transcriptional start site is indicated with an arrow, introns are shown as black lines and exons as coloured blocks. (**b**) Quantifications of mesoblast lineage descendants in the indicated genotypes, in the tail area (early dividing body wall muscles) and around the vulva (late dividing egg-laying muscles).

### The SWI/SNF cell-cycle arrest function is dosage sensitive

We considered whether different levels of SWI/SNF BAF may explain the opposite over-proliferation and proliferation-arrest phenotypes. Dosage sensitivity of SWI/SNF complex functions has been implied by the spectrum of mutations detected in human cancer and intellectual disability disorders (Kadoch et al. 2013; Sun et al. 2017; Kosho et al. 2014; Bögershausen and Wollnik 2018). In both, heterozygous loss-of-function mutations in SWI/SNF BAF subunit encoding genes are common, revealing haploinsufficiency or dominant-negative effects (Kadoch and Crabtree 2015; Wilson and Roberts 2011). Moreover, specific cancer cells deficient for the BRG1 ATPase or ARID1A display synthetic lethal interactions with the BRM ATPase and ARID1B subunits, respectively, which have partly overlapping functions (Helming et al. 2014b; Hoffman et al. 2014; Wilson et al. 2014). Thus, at least in some cancers, SWI/SNF subunit genes behave as dominant tumor suppressors, while in several instances full SWI/SNF BAF inactivation is incompatible with cancer cell viability. In our *C. elegans* model, the heterozygous *swsn-4*^KA^/wt BRG1/BRM mutation did not alter the mesoblast proliferation-differentiation pattern (Figure 4a). However, when combined with *lin-23* RNAi or a *cki-1*^Lox^ conditional knockout, a single inactive *swsn-4* allele (*swsn-4*^KA^/wt) led to a substantial increase in the number of M cell descendants (Figure 4a). Thus, our *in vivo* observations in a reproducible lineage support that the cell-cycle arrest function of SWI/SNF ATPases is dosage sensitive and probably haploinsufficient.

**Figure 4.**
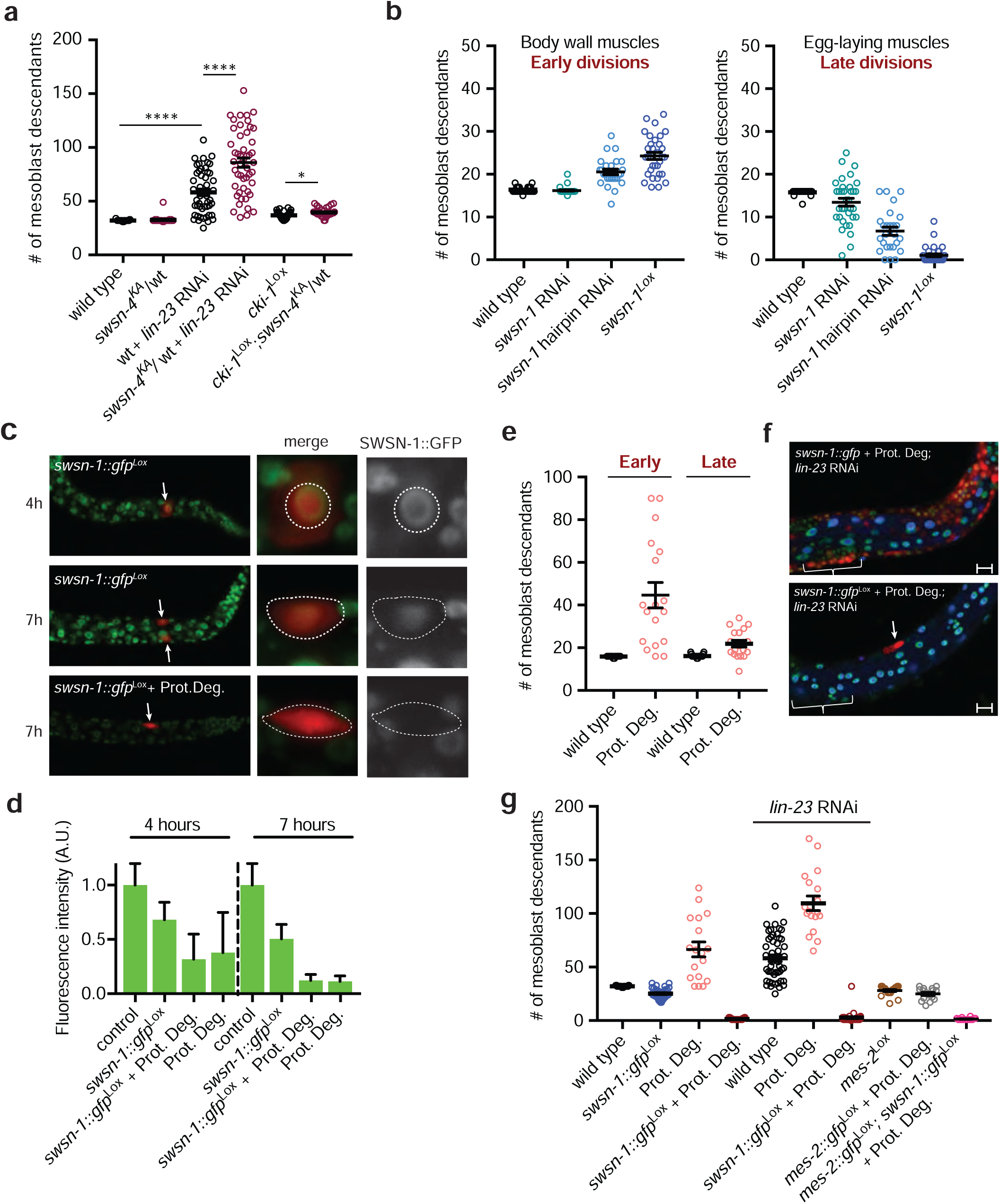
Opposite and acute cell division phenotypes depending on the dosage of residual SWI/SNF activity. (**a**) Quantification of total number of mesoblast descendants for the indicated genotypes. SWI/SNF haploinsufficiency is visible when a heterozygous null allele is combined with loss of a negative cell cycle regulator *lin-23*^ßTrCP^ or *cki-1*^Kip1^. (**b**) Quantification of mesoblast descendants for the indicated genotypes in the tail area (early divisions) and around the vulva (late divisions). Expression of a double stranded hairpin RNA against *swsn-1*, under the control of P*hlh-8*, leads to stronger loss of function than L1 feeding RNAi and a phenotype resembling gene knockout. (**c**) Expression of SWSN-1::GFP in the M cell at 4 and 7 hours of larval development following gene knockout, or gene knockout combined with protein degradation (Prot. Deg., bottom). Arrows indicate mesoblast cells in the larva views, which are outlined in the zoom images. (**d**) Quantification of SWSN-1::GFP by fluorescence intensity in M at the indicated times and genotypes. (**e**) Quantifications of mesoblast descendants for the indicated genotypes, in the tail area (early) and around the vulva (late). Note that SWSN-1::GFP degradation alone (Prot. Deg.) leads to overproliferation of both early and late dividing mesoblast descendants. (**f**) Representative images of tumourous overproliferation following SWSN-1 protein degradation and *lin-23* RNAi (top), and of the one-cell arrest (arrow) after *swsn-1* gene knockout together with protein degradation in *lin-23* knockdown conditions (bottom). The brackets indicate the vulva region. Scale bar 10 µm. (**g**) Quantification of total number of mesoblast descendants for the indicated genotypes. Protein degradation alone of SWSN-1::GFP (Prot. Deg.) leads to overproliferation, which is strongly increased by RNAi of *lin-23*, whereas protein degradation combined with gene knockout (*swsn-1*::*gfp*Lox + Prot. Deg.) leads to acute cell division arrest. RNAi of *lin-23* and acute knockout of EZH2-related *mes-2* (*mes-2*::*gfp*Lox + Prot. Deg.) do not suppress the acute one-cell arrest phenotype.

### A low level of SWI/SNF BAF is required for cell proliferation

More effectively than missing one allele of *swsn-4*, extra cell divisions result from RNAi of SWI/SNF BAF subunits, the *swsn-1(ts)* mutation (Ruijtenberg and van den Heuvel 2015), and - initially-SWI/SNF gene knockout. We hypothesized that in each of these situations, hyperplasia is associated with incomplete loss of SWI/SNF function. Following CRE-Lox mediated gene excision, residual mRNA and protein will initially remain present, which will be depleted with time and additional cell divisions. This means that the appearance of a true null phenotype will be delayed with respect to the timing of gene excision, and could be manifested as cell division arrest. To test this hypothesis, we sought to better control SWI/SNF inactivation. First, we started RNAi earlier, by expressing a *swsn-1* dsRNA hairpin in the embryonic mesoblast, controlled by the *hlh-8* Twist promoter. This lineage-specific RNAi created a phenotype in between that observed after RNAi by feeding L1 larvae and gene knockout, inducing overproliferation of the early BWM precursors as well as cell division arrest of the late-dividing egg-laying muscle precursors (Figure 4b).

To further examine and control the degree of SWI/SNF gene inactivation, we made use of the combined insertion of a GFP tag and Lox sites into the endogenous *swsn-1* gene (Figure 2a). Following lineage-specific gene knockout, SWSN-1::GFP expression was still detectable in the mesoblast before (4h) and at the time (7h) of the first post-embryonic cell division, although at progressively lower levels as compared to the wild type (Figure 4c-d). These observations support that despite SWI/SNF gene knockout, residual protein remains present during the early mesoblast divisions.

To create an acute null phenotype, we combined the conditional gene knockout with lineage-specific protein degradation. To achieve this, we expressed an anti-GFP-nanobody::ZIF-1 fusion protein in the mesoblast lineage (Wang et al. 2015, 2017). The fusion protein targets GFP to a CUL-2 based E3 ubiquitin ligase, thereby triggering efficient SWSN-1::GFP proteolysis. We integrated an inducible transgene, which upon CRE-mediated removal of a STOP cassette expresses anti-GFP nanobody::ZIF-1 from the ubiquitous *eft-3* promoter (figure supplement 5a-b). This way, CRE expression in the mesoblast induces protein degradation in parallel to *swsn-1::gfp* gene excision. The double inactivation approach reduced SWSN-1::GFP expression to undetectable levels before the first larval division (Figure 4d). In fact, nanobody-mediated degradation alone, without gene knockout, also led to strong SWSN-1::GFP depletion, but in this setting GFP remained detectable at later time points (Figure 4d-f).

Strikingly, the effects of SWSN-1::GFP degradation alone versus degradation plus gene knockout were completely opposite. Lineage-specific protein degradation resulted in the strong overproliferation of early as well as late muscle precursors cells (Figure 4e). The overproliferation of both early and late precursors was further enhanced by simultaneous knockdown of a negative cell cycle regulator (*lin-23*; Figure 4f-g). Thus, the low residual levels of SWSN-1::GFP in M descendants after GFP-directed protein degradation are sufficient to sustain proliferation, but not to induce timely cell-cycle arrest. By contrast, simultaneous SWSN-1::GFP protein degradation and gene knockout resulted in a complete block of cell division of the mesoblast precursor cell (Figure 4g). Even when combined with *lin-23* knockdown, the single embryonic mesoblast did not enter cell division during larval development in most animals (Figure 4f). These data clearly demonstrate that the SWI/SNF complex functions are dosage sensitive, with a complete absence being incompatible with proliferation, while reduced levels promote overproliferation.

### An essential SWI/SNF complex function, independently of PcG protein opposition

To test whether unopposed PcG activity contributes to the mesoblast arrest, we adjusted the endogenous *mes-*2 H3K27 methyltransferase locus for conditional complete inactivation. Gene excision of *mes-2::gfp*^Lox^ combined with anti-GFP nanobody::ZIF-1 expression did not interfere with early M divisions, although premature arrest of cell division - and possibly initiation of differentiation - occurred late in the lineage (figure supplement 5c). Acute double inactivation of MES-2 and SWSN-1 showed the single-cell arrest phenotype, indicating that PcG PRC2 loss does not alleviate the SWI/SNF requirement (Figure 4g). These data support that SWI/SNF complexes are required for an essential function beyond opposing PcG-mediated gene repression.

As expected for PcG-mediated transcriptional repression, the effects of *mes-*2 PRC2 inactivation became apparent only after four to seven rounds of cell division. This is in stark contrast to the immediate arrest of SWI/SNF null mesoblasts. As the arrested cells remained present even in old adults, the SWI/SNF BAF requirement may reflect a function for the complex in cell proliferation.

### Acute SWI/SNF depletion leads to cell division arrest before DNA replication

Quantitative measurements of the DNA content of arrested SWI/SNF null mesoblasts demonstrated that these cells stopped the cell cycle before S, or in very early S phase (Figure 5a). Therefore, it is highly unlikely that the arrest results from a TOP2A-associated function of BAF complexes, which has been reported to contribute to faithful mitosis (Dykhuizen et al. 2013). We considered the possibility of a DNA damage or intra-S-phase checkpoint arrest, as SWI/SNF complexes have also been implicated in DNA damage repair and replication (Kadoch and Crabtree 2015; Brownlee et al. 2015). To alleviate DNA damage and S phase checkpoints, we added high concentrations of exogenous dNTPs, and performed RNAi of *chk-1*, and double RNAi of *lin-35* Rb and *cep-1* p53 (Kalogeropoulos et al. 2004; Bester et al. 2011). As none of these conditions affected the single-cell arrest phenotype, evidence for checkpoint arrest was not obtained (Figure 5b).

**Figure 5.**
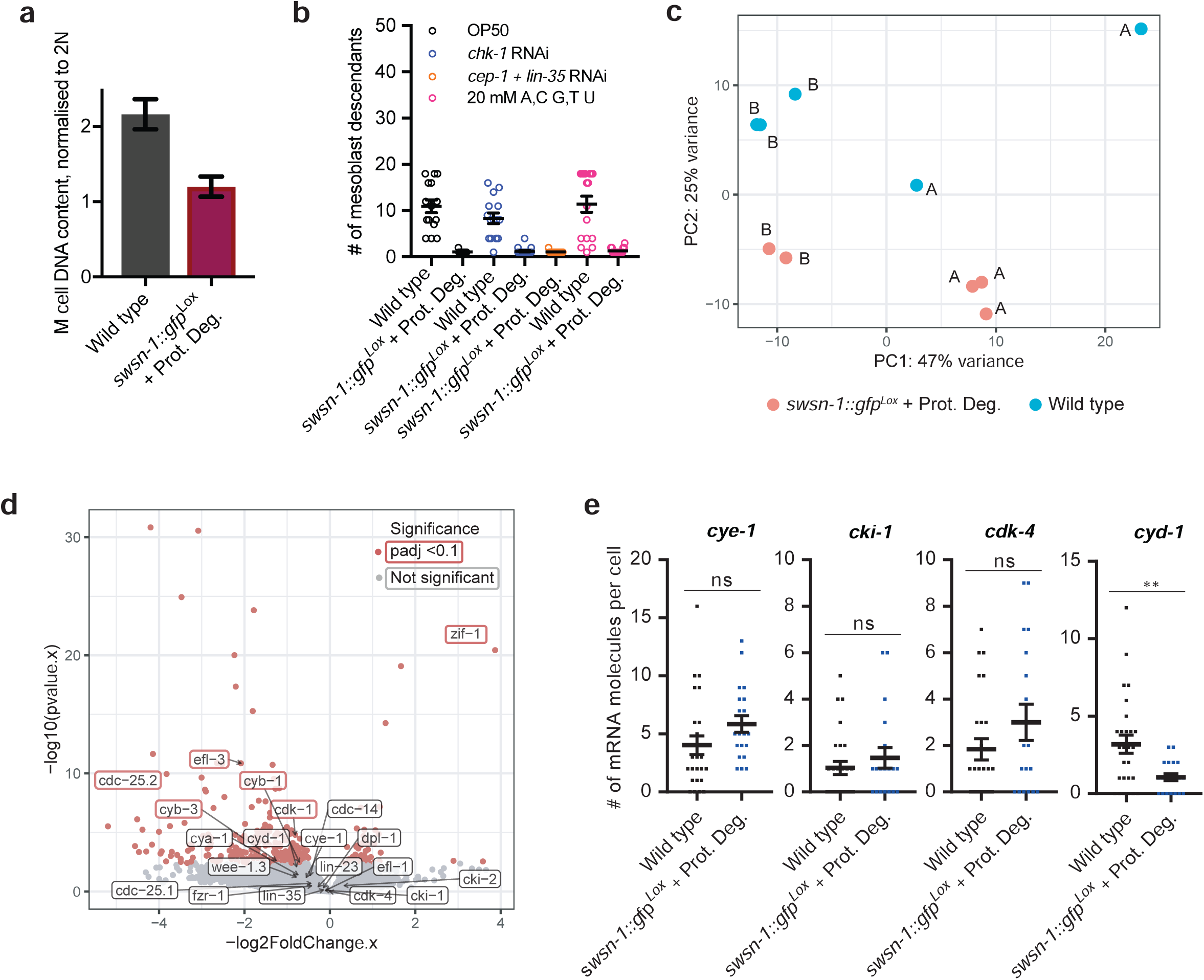
Transcriptome RNA-sequencing demonstrates a continuous requirement for SWI/SNF BAF complexes in normal transcription and proliferation control. (**a**) Quantification of the DNA content of the M cell in synchronised L1 larvae after 7 hours of larval development of animals with the indicated genotypes. Note that wild type M cells have undergone S phase but not yet divided, leading to a 4N DNA content, whereas M cells in acute *swsn-1::gfp* knockout animals show an approximately 2N DNA content, indicative of cells arresting in G1 or early S phase. DNA was stained using propidium iodide, and DNA content quantified in ImageJ, normalised to the average of differentiated embryonically formed body wall muscle cells (2N). (**b**) Quantifications of mesoblast descendants for the indicated genotypes and treatments at 12 hours of development. (**c**) Principal component analysis indicating clustering of replicate RNA-sequencing libraries, prepared from FACS sorted 2000-cell samples from wild type and *swsn-1::gfp*^Lox^ + Protein Degradation L1 larvae at 5.5 hours of development. Samples “A” and “B” are true biological replicates, with RNA-sequencing libraries prepared from different starting populations of synchronised worms, on different dates. Within A and B, duplicate/triplicate RNA-sequencing libraries were prepared from different 2000-cell populations, isolated from the same starting worm population, and can thus be considered “semi-biological” replicates (**d**) Volcano plot indicating differentially expressed genes between *swsn-1:gfp*^Lox^ + Protein Degradation and wild type isolated mesoblast cells, at 5.5 hours of development. Differential expression analysis was carried out using DEseq2, with 5 experimental replicates for each strain. (**e**) Quantifications of the number of mRNA molecules per cell in smFISH experiments for the indicated genes, in synchronised L1 larvae at 6.5 hours of larval development (just before the usual time of the first M division), in wild type compared to *swsn-1::gfp*^*Lox*^ + Protein Degradation.

### The SWI/SNF complex is continuously required and promotes expression of cyd-1 cyclin D1

We performed whole transcriptome RNA-sequencing (RNA-Seq) to further characterize the arrested cells. We used pools of 2000 wild type or acute *swsn-1* knockout mesoblasts, isolated from synchronous cultures of L1 larvae at 5.5 hours of development (1-1,5 hours before the normal time of the first mitosis). Principal component analysis (PCA) showed a clear separation of the wild type and mutant sequence data sets (Figure 5c). Nevertheless, only a limited number of genes showed significantly different expression, of which the large majority were reduced in the SWSN-1 depleted mesoblasts (213 genes; Supplementary Table 2). Among those, cell-cycle genes were well represented, in particular presumed E2F targets (e.g. *cdc-25.2, cdk-1, cyb-1* cyclin B1, and *cyb-3* cyclin B3) (Figure 5d, red boxes). These genes encode regulators of the G2/M transition and are expected to be expressed in wild type cells, which at this stage are preparing for mitosis, but not in G1-arrested cells. Therefore, the reduced transcript levels of these cell-cycle genes may result indirectly from the early cell-cycle arrest of *swsn-1* mutant mesoblasts. Regulators of the G1/S transition, such as *cdk-4* CDK4/6, *cye-1* cyclin E, *cki-1* p21 and *lin-35* Rb, showed similar expression in wild type and arrested mesoblasts (Figure 5d, grey boxes). As a possible exception, the reduced number of *cyd-1* cyclin D transcripts in *swsn-1* mutant mesoblasts was significant in one of the two biological replicates. As cyclin D transcription is an important regulator of cell-cycle entry, we followed up on this finding by examining transcript numbers with single-cell resolution. Using smFISH, we observed a significantly lower number of *cyd-1* mRNA molecules in *swsn-1* mutant mesoblasts compared to normal mesoblasts, even before the first cell division (6,5 hrs; Figure 5e). These data demonstrate a continuous requirement for SWI/SNF BAF complexes in normal transcription and proliferation control, even when PcG-mediated repression is severely reduced.

## Discussion

In this study, we examined SWI/SNF and PcG complex functions in an *in vivo* system that provides a well-defined cellular context and reproducible developmental decisions. Our gene knockdown and knockout experiments demonstrate that in the same cell type and conditions, reducing the level of SWI/SNF core subunits or ARID1 factor interferes with cell-cycle withdrawal and the transition to a more differentiated state, while complete inactivation of the identical subunits is incompatible with cell proliferation. Together with EZH2-knockout studies, our data imply that the SWI/SNF BAF ATPase exerts a tumor suppressor function in the mesoblast lineage that requires a relatively high functional level and involves PcG opposition, while a low level is essential and sufficient to sustain cell proliferation. Our data are consistent with the model that partial loss-of-function mutations are selected during carcinogenesis because they reduce a differentiation-promoting tumor suppressor activity without inactivating the critical requirement for the SWI/SNF complex.

That the function of SWI/SNF complexes is dosage sensitive was previously concluded from the heterozygous and sometimes subtle mutations in SWI/SNF subunits identified in human cancer and neurologic disease (Kadoch and Crabtree 2015; Bögershausen and Wollnik 2018). The finding that incomplete inactivation of multiple different BAF subunits resulted in a similar hyperplasia phenotype indicates that the reduced SWI/SNF function, rather than the activity of complexes with an abnormal subunit composition, lead to overproliferation in our system. Hyperplasia was observed early and reproducibly after SWI/SNF BAF gene knockout or protein degradation. This implies that neither a dominant-negative mechanism, nor the generation of secondary mutations, as a result of DNA damage or defective TopoII function (Kadoch and Crabtree 2015), are needed for a hyperplastic response.

The situation in human cancer is obviously much more complex and includes various mechanisms of SWI/SNF deregulation. As such, common heterozygous mutations in the BRG1 ATPase selected in human cancer are likely to act in a dominant-negative way (Hodges et al. 2018). Nevertheless, such mutations do not fully inactivate the wild type allele, and residual BRG1 function is thus retained in the heterozygotes. This is in agreement with the essential SWI/SNF function detected in our system, as well as in cancer cells by synthetic lethal screening (Helming et al. 2014b; Hoffman et al. 2014; Wilson et al. 2014). However, some cells appear to survive without SWI/SNF function. In a specific small cell carcinoma of the ovary as well as a subset of cancer-derived cell lines, biallelic mutation of BRG1 coincides with transcriptional silencing of the BRM locus (Kadoch and Crabtree 2015; Shain and Pollack 2013; Karnezis et al. 2016). In addition, malignant rhabdoid tumors (MRTs) are well-known for their homozygous loss of SMARCB1 (SNF5), one of the core subunits of SWI/SNF complexes. Although this was initially expected to fully disable SWI/SNF complexes, the growth of SNF5-deficient tumors still requires BRG1 (Wang et al. 2009). Recent studies revealed the presence of ncBAF complexes that do not contain SMARCB1, while removal of ncBAF-specific subunits induces synthetic lethality in cancer cells lacking SMARCB1 (Mashtalir et al. 2018; Michel et al. 2018). Thus, although compensatory mechanisms may be present or selected for in some cell types, the complete loss of all SWI/SNF ATPase activity is in general incompatible with cell proliferation.

Several mechanisms could underlie the dosage sensitivity of SWI/SNF functions. A high dosage appears needed to alter the expression of many loci when cells transition to the differentiated state. If the maintenance of transcription can be achieved with much lower levels of the SWI/SNF complex, partial SWI/SNF inactivation will interfere with differentiation but not continued proliferation, which provides a mechanism promoting tumorigenesis. Such a mechanism is supported by the alterations induced by SMARCB1 or SMARCA4 knockout in mouse embryonic fibroblasts (MEF), which most significantly reduced the number of transcripts from genes with GO terms associated with development and differentiation (Alver et al. 2017).

The knockout of SWI/SNF subunits in MEF cells, as well as introduction of heterozygous dominant-negative alleles of BRG1 in mouse embryonic stem cells caused a broad reduction of chromatin accessibility at active enhancers, which remarkably was associated with loss of H3K27Ac rather than increased PcG protein binding (Hodges et al. 2018; Alver et al. 2017). This does not reveal whether widespread transcriptional deregulation, reduced expression of some critical genes, or other defects are incompatible with cellular proliferation when the SWI/SNF activity falls below a critical level. Our RNA-seq analysis showed that the acute arrest of mesoblast proliferation occurred when the expression of a limited number of genes was significantly altered. We identified *C. elegans* cyclin D as one of the genes whose transcription is acutely sensitive to SWI/SNF inactivation. The proliferation arrest associated with strong SWI/SNF loss was insensitive to knockdown of cell cycle inhibitors, however, in contrast to *cyd-1* mutants (Boxem et al. 1999; The et al. 2015). Therefore, we do not expect that the downregulation of *cyd-1* alone is responsible for the tight cell-cycle arrest. Remarkably, two recent human cancer studies also concluded that SWI/SNF ATPases promote cyclin D1 expression (Xue et al. 2019b, 2019a). This was found to underlie a synthetic lethal interaction between CDK4/6 inhibition and SMARCA4 loss in SCCOHT ovarian carcinoma and non-small cell lung cancers.

That loss-of-function of SWI/SNF subunits can lead to opposite phenotypes (hyperplasia versus division arrest), depending on the residual levels, provides support for the clinical exploration of cancer cell vulnerabilities that result from SWI/SNF gene mutations (Helming et al. 2014a; St Pierre and Kadoch 2017). At the same time, the delicate balance between dosage-dependent SWI/SNF and PcG regulators observed in our system illustrates that the outcome of targeted therapies will be difficult to predict and highly context dependent. The many parallels between observations in our system and human cancer cells supports taking advantage of efficient genetic screening in *C. elegans* to identify synthetic lethal interactions that are broadly associated with SWI/SNF loss and cause little toxicity in normal cells.

## Materials and methods

### Strains and culture

Genotypes of all strains used in this study are listed in Supplementary Table 1. *C. elegans* was cultured on NGM plates seeded with OP50 bacteria and generally maintained at 20°C. Strains containing the *pha-1(e2123)* mutation were maintained at 15°C and shifted to 25°C for mutant phenotype analysis.

### RNA-mediated interference

Bacterial cultures of *E. coli* HT115 containing L4440 empty vector or vector with genomic or ORF gene inserts were grown overnight, induced with 1 mM IPTG for 1 hour, 2.5 times concentrated and seeded onto NGM plates containing 12,5 μg/ml Tetracycline, 100 μg/ml Ampicillin and 2 mM IPTG. RNAi feeding for knock-down of SWI/SNF components initiated from L1 stage and the number of mesoblast descendants was analyzed in late L3/early L4 animals of the same generation. RNAi clones were obtained from the Vidal (Lamesch et al. 2004) and Ahringer (Fraser et al. 2000) databases and sequence verified. *swsn-1, swsn-4, swsn-8, swsn-7, swsn-9, swsn-2.2, swsn-3, swsn-6* and *phf-10* RNAi vectors were cloned by ligating a ∼1000 bp cDNA fragment into the L4440 vector. A different RNAi approach was used in order to analyze the role of *lin-23, chk-1, lin-35* and *cep-1*. L4 animals were placed on RNAi plates and the F1 was analyzed.

### Molecular cloning

For the MosSCI recombination reporter (readout^lox^), P*eft-3*::LoxP::*egl-13* NLS::tagBFP2::*tbb-2* UTR::LoxP::*egl-13* NLS::mCherry::*tbb-2* UTR, two separate Gibson assembly reactions were performed to generate readout^lox^ part one (P*eft-3*::LoxP::*egl-13* NLS::tagBFP2::*tbb-2* UTR) and readout^lox^ part two (LoxP::*egl-13* NLS::mCherry::*tbb-2* UTR) in pBSK vectors. Next, a third Gibson assembly reaction was performed to introduce both readout^lox^ parts into the same pBSK vector. Finally, the 9349 bp readout^lox^ cassette was cloned as a MluI and NotI fragment into the pCFJ350 MosSCI vector. See figure supplement S5 for details on the anti-GFP nanobody-ZIF-1 construct. The *myo-3::H2B::GFP* was previously used in (Korzelius et al. 2011). For the generation of the MosSCI construct P*hlh8*::GFP::H2B::*unc-54* UTR, a PCR product containing GFP::H2B::*unc-54* UTR was obtained using the P*wrt-2*::GFP::H2B (Wildwater et al. 2011) construct as a template. In addition, a PCR fragment of 518 bp directly upstream of the *hlh-8* ATG was obtained by using P*hlh-8*::CRE (Ruijtenberg and van den Heuvel 2015) as a template. Next, the PCR fragments were cloned into the pCFJ350 MosSCI vector using Gibson assembly. The P*hlh8*::*swsn-1*::*unc-54* UTR hairpin construct was generated by replacing GFP::H2B in the above mentioned P*hlh-8* containing vector with a ∼1100 bp *swsn-1* cDNA Mlu/MscI fragment. Next, the antisense sequence of the same ∼1100 bp *swsn-1* cDNA fragment was cloned using primers containing the restriction sites MscI and NheI. The NheI containing primer also introduced a stop codon.

### CRISPR/Cas9 genome editing

All Lox insertions, as well as the ATPase dead *swsn-4* mutant, were generated by temperature sensitive *pha-1* co-conversion (Ward 2015) using single-stranded DNA oligonucleotides with 40 bp homology arms as repair templates. *pha-1* co-conversion was performed as follows. Young, 7 times out-crossed, *pha-1(e2123)* adults grown at the permissive temperature of 15°C were injected into the gonads with the following injection mix: 50 ng/µl U6::gRNA target construct, 60 ng/µl pJW1285 (U6::gRNA *pha-1* and Cas9 construct) and 50 ng/µl ssDNA repair templates for *pha-1* as well as the appropriate target. Injected worms were immediately placed at 25°C. F1 *pha-1* rescue animals were analyzed by PCR using primers flanking the Lox insertion site showing a band shift upon Lox insertion. The *swsn-4* K to A mutation was generated in a strain which has an integrated cassette, *eft-3*::GFP::2xNLS::*tbb-2* 3’UTR, on chromosome IV close to the K to A mutation site. By selecting GFP positive worms, heterozygous ATPase-dead mutant worms can be maintained. The K to A mutation was identified using a primer annealing to the mutated sequence. All mutations were sequence verified. The *swsn-1*::*egfp*^LoxP^ strain was generated by inserting eGFP using the self-excising cassette (SEC) method (Dickinson et al. 2015) into wild type N2 worms, which leaves a LoxP scar. The injection mix to generate the GFP for the creation of the *swsn-1*::*egfp*^LoxP^ insertion contained: 100 ng/µl U6::gRNA target construct, 20 ng/µl pDD268 eGFP SEC vector with 150 bp *swsn-1* Left Homology arm and *swsn-1* 600 bp Right Homology arm, 50 ng/µl P*eft-3*::Cas9 (Addgene 46168) and 2,5 ng/µl *Pmyo-2*::*tdTomato.* The second LoxP site was introduced by crossing the strain with *pha-1*(*e2123*) and by temperature-sensitive *pha-1* co-conversion as described above. The *mes-2::egfp*^LoxN^ strain was made by inserting eGFP containing a LoxN site (ordered as a gBlock from IDT) in one of the GFP introns, into a strain which already contained a LoxN site in the first intron of *mes-2*.

### Microscopy and quantification of cell numbers

Expression of *hlh-8::*CRE in combination with the lineage-tracing reporter allowed us to specifically visualize the mesoblast lineage (mCherry-positive). Extra division events were quantified by counting the number of mCherry-positive cells. Images of the overproliferation phenotypes were obtained using a Zeiss LSM700 Confocal microscope. Quantification of SWSN-1::GFP fluorescence intensity was done using the ImageJ measurement tool, by selecting the region of interest (M cell) and subtracting background signal. At least 16 larvae per condition were measured.

### Single-molecule fluorescence in situ hybridization (smFISH)

smFISH was performed essentially as described in (Ruijtenberg and van den Heuvel 2015). Cy5-coupled probes against mRNA’s of interest were ordered from Stellaris (http://singlemoleculefish.com/), with 23-48 probes ordered per gene of interest ranging from 18-22 bp in length. L1 or L4 worms were fixed using Bouins fixative as previously described (van den Heuvel and Kipreos 2012). In short: animals were fixed for 30 min. at RT in 400 μl Bouin’s fix + 400 μl methanol and 10 μl β-mercaptoethanol, three times freeze-thawed and again tumbled for 30 min in fixative at RT. For permeabilization, the fixative was removed and exchanged for borate-Triton-β-mercaptoethanol (BTB: 1xBorate Buffer, 0.5% Triton and 2% β-mercaptoethanol) solution. Animals were tumbled 3 times for 1 hour in BTB solution at RT. BTB solution was then replaced with wash buffer A (Stellaris) containing 20% formamide, and then with 100 µl hybridization solution containing smFISH probes to a final concentration of 0.25-0.5 µm, and incubated overnight at 32 °C. Samples were washed in Wash buffer A without formamide, and incubated with 0.05 ng DAPI in wash buffer A for 30 minutes at 32 °C. After a final wash in wash buffer B (Stellaris), worms were mounted on slides with Vectashield mounting medium and imaged within 4 hours. Images were acquired using a Nikon Eclipse Ti Spinning Disk Confocal microscope, using the 100x objective. The TMR (mCherry) spots were used to draw a region of interest around (mCherry-positive) M lineage descendants in ImageJ, in which the number of Cy5 fluorescent mRNA spots was quantified using the ComDet plugin in ImageJ (https://imagej.net/Spots_colocalization_(ComDet)).

### Statistical analysis

Sample sizes were not pre-determined; instead, all available animals of the right stage and genotype were counted. smFISH data are included from at least eight independent animals, reporter expression and cell numbers from at least ten independent animals. Graphs and data analysis were produced using GraphPad Prism 6.05. Plots indicate all data points, as well as the mean (average) ± SEM. As the data essentially fit normal distributions, unpaired two-tailed Student’s t tests were used to examine statistical significance of the difference between means.

### Propidium iodide staining

Propidium iodide staining was carried out after Carnoys fixation as previously described (Boxem et al. 1999). For DNA quantification, Z-stacks were acquired using a Zeiss LSM700 confocal microscope. Maximum projections (SUM) were made in ImageJ from all the stacks in which the M cell DNA was visible, and pixels quantified using ImageJ. Post-mitotic, differentiated body wall muscle cells (2N) were quantified in the same manner and used as a reference.

### Larval cell isolation

The *C. elegans* L1 larval cell isolation protocol was adapted from (Zhang et al. 2011). To generate large amounts of synchronized L1 larvae, worms were grown in S medium in liquid culture for two generations (to enrich for gravid adults) and bleached. Eggs were hatched overnight (for 18-22 hours) in S medium without food, and starved L1 larvae split into three aliquots and put back into S medium with OP50 for 5.5 hours. Cultures were put on ice for 15 minutes, spun down at 1300g, washed 2x in M9 and once in H20. L1 larvae were then transferred to 1.5 ml Eppendorf tubes (20-40 µl L1 pellet per Eppendorf) and spun down at 16000g. Larvae were treated with SDS-DTT solution (20 mM HEPES pH 8.0, 0.25% SDS, 200 mM DTT, 3% sucrose) for 2 mins, washed 6 times in egg buffer (25 mM HEPES pH 7.3, 118 mM NaCl, 48 mM KCl, 2 mM CaCl2, 2 mM MgCl2, adjusted to 340±5 mOsm with H20) then treated with 20 mg/ml Pronase E in L15/FBS buffer (10% FBS and 1% Penicillin-Streptomycin -Sigma P4458-in L-15 insect medium, adjusted to 340±5 mOsm with 60% sucrose), for 30-40 minutes. After 10 minutes, and 20 minutes, in Pronase E, a pellet pestle motor with a pellet pestle adapted to 1.5 ml microtubes (Sigma Z359971 and Z359947) was used for 1 min on each sample. Finally, cell preparations were washed 3× in L15/FBS, spun down at 9600g for 5 minutes between each wash, and resuspended in 1 ml L15/FBS.

### FACS sorting

Cell preparations were allowed to settle on ice for 30 minutes, and the top 850 µls of supernatant was removed and transferred to a new Eppendorf tube for FACS sorting. Cells were sorted according to mCherry positive signal using a BD FACSARIA III (BD Biosciences). For each sample 2000 cells were sorted into L15/FBS buffer. In one session, three wild type and three mutant samples were sorted. Immediately after sorting, cells were spun down at 12000 g for 5 minutes, resuspended in TRIzol and frozen at −80°C.

### cDNA library preparation

cDNA libraries were prepared according to a combination of the CEL-Seq and CEL-Seq2 protocols with some modifications (Hashimshony et al. 2012, 2016). RNA was precipitated using chloroform/isopropanol precipitation at −20°C for 48-72 hours, and washed once in 75% ethanol. CEL-Seq2 primers were used (one unique primer per sample), with each primer containing an anchored polyT, a 6 bp unique barcode, 6 bp UMI, a 5’ Illumina adaptor and a T7 promoter. The CEL-Seq1 protocol was followed for a first round of reverse transcription and cDNA cleanup followed by in vitro transcription, as well as for fragmentation of amplified RNA (aRNA), as described (Hashimshony et al. 2012). aRNA was run on an agilent bioanalyzer (RNA pico-ChIP) for quality control and quantification. The CEL-Seq2 protocol was followed for a second round of reverse transcription and PCR amplification, as described (Hashimshony et al. 2016). cDNA was amplified for 11-15 cycles depending on aRNA amounts, run on an agilent bioanalyzer (DNA picoChIP), quantified using a Qubit and 1-2 ng sequenced with 5% coverage on an Illumina NextSeq500.

### RNA-sequencing data analysis

Data analysis was carried out in R version 3.4.4. Principal component analysis (PCA) was performed with the plotPCA function after carrying out variance-stabilized transformation on the data. Differential gene expression was analyzed using DESeq2 (default settings) using a padj cutoff of 0.1 (Love et al. 2014). Plots were generated using ggplot2 (Wickham 2016).

## Supporting information

supplemental figures and tables van der Vaart et al

## Acknowledgements

We are grateful to members of the Korswagen and Van Oudenaarden groups at the Hubrecht Institute for help with RNA-Seq experiments, and to P. Verrijzer and members of the Van den Heuvel and Boxem groups for input, discussion and comments on the manuscript. Several strains were provided by the CGC, which is funded by the NIH National Center for Research Resources (NCRR).

## Author contributions

AvdV performed the majority of the SWI/SNF knockdown and knockout experiments, analyzed the results and co-wrote the paper. MG generated *cki-1*^Lox^ and *mes-2::gfp*^*Lox*^ strains, performed the smFISH, DNA quantification and RNA-Seq experiments, analyzed results and co-wrote the paper. VP generated the anti-GFP nanobody::ZIF-1 construct and strains and contributed to experimental design. SvdH conceived the study, acquired funding, provided guidance with experimental design and co-wrote the manuscript.

