## supplemental figures and tables van der Vaart et al for "Dose-dependent functions of SWI/SNF BAF in permitting and inhibiting cell proliferation *in vivo*"

**a**

| *C. elegans* gene | alternative *C. elegans* name | mammalian homologue(s) |
|---|---|---|
| *swsn-1* | *psa-1* | SMARCC1/BAF155; SMARCC2/BAF170 |
| *swsn-4* | *psa-4* | SMARCA2/BRM; SMARCA4/BRG1 |
| *snfc-5* | *swsn-5* | SMARCB1/BAF47/SNF5 |
| *swsn-8* | *let-526, psa-10, let-104, lss-4* | ARID1/BAF250 |
| *pbrm-1* | *tag-185* | PBRM1/BAF180 |
| *swsn-7* | | ARID2/BAF200 |
| *swsn-9* | *tag-298* | BRD7/BRD9 |
| *swsn-2.1* | *ham-3* | SMARCD/BAF60 |
| *swsn-2.2* | | SMARCD/BAF60 |
| *swsn-3* | | SMARCE1/BAF57 |
| *swsn-6* | *psa-16* | ACTL6/BAF53 |
| *bcl-7* | | BCL7 |
| *dpff-1* | | DPF/BAF45 |
| *phf-10* | | PHF10/BAF45 |

**b**

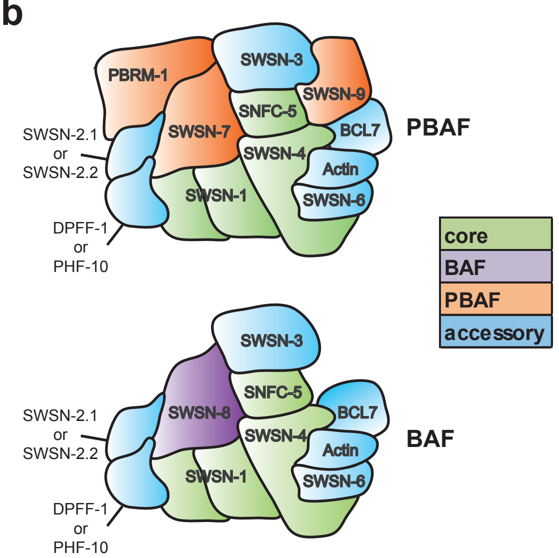


**c**

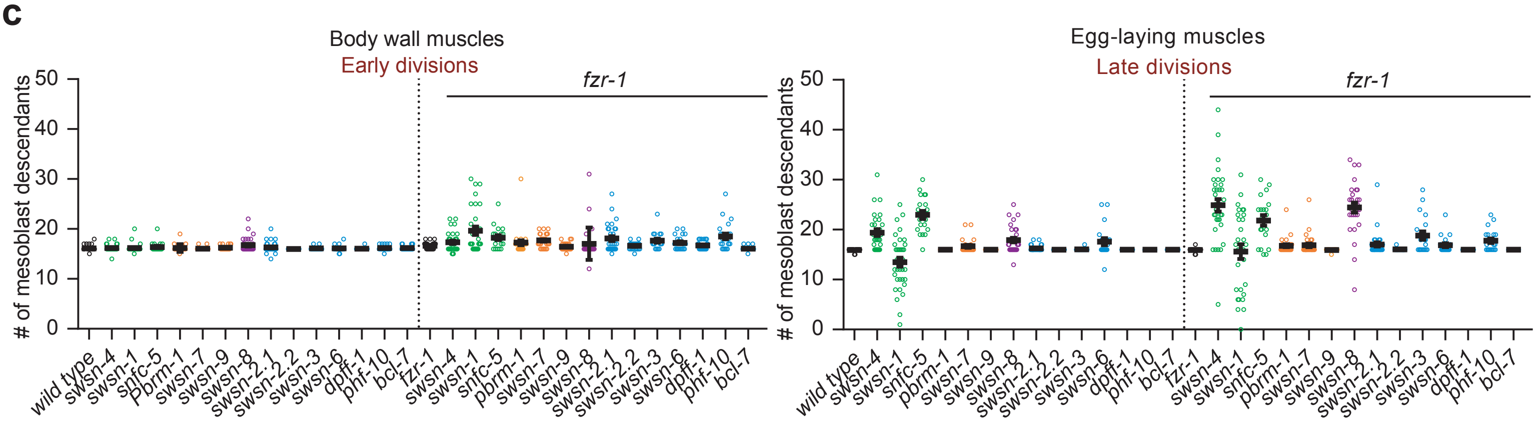


**d**

| | *# cells in the tail (Early muscle)* | | | | *# cells around the vulva (Late muscle)* | | | | *total # of cells* | | | |
|---|---|---|---|---|---|---|---|---|---|---|---|---|
| gene | wild type | sig. | *fzr-1* | sig. | wild type | sig. | *fzr-1* | sig. | wild type | sig. | *fzr-1* | sig. |
| L4440 | 16.13 ± 0.56 | | 16.64 ± 0.70 | | 15.94 ± 0.25 | | 15.96 ± 0.35 | | 32.06 ± 0.63 | | 32.60 ± 0.76 | |
| *swsn-1* | 16.17 ± 0.79 | ns | 19.67 ± 4.58 | ** | 13.49 ± 5.39 | * | 15.60 ± 8.15 | ns | 29.66 ± 5.15 | * | 35.27 ± 5.47 | * |
| *swsn-4* | 16.20 ± 0.71 | ns | 17.27 ± 1.98 | ns | 19.40 ± 3.69 | **** | 25.80 ± 7.09 | **** | 35.60 ± 3.29 | **** | 43.07 ± 6.47 | **** |
| *snfc-5* | 16.40 ± 0.94 | ns | 18.25 ± 2.20 | ** | 22.95 ± 3.68 | **** | 21.85 ± 4.74 | ns | 39.35 ± 3.66 | **** | 40.10 ± 5.94 | **** |
| *swsn-8* | 16.75 ± 1.41 | * | 17.03 ± 3.23 | ns | 17.91 ± 2.63 | *** | 24.43 ± 5.33 | **** | 34.66 ± 2.67 | **** | 41.47 ± 4.31 | **** |
| *pbrm-1* | 16.19 ± 0.75 | ns | 17.25 ± 2.58 | ns | 16.00 ± 0.00 | ns | 16.79 ± 1.62 | * | 32.19 ± 0.75 | ns | 34.04 ± 2.94 | * |
| *swsn-7* | 16.08 ± 0.28 | ns | 17.70 ± 1.26 | *** | 16.67 ± 1.47 | ** | 16.87 ± 2.01 | * | 32.75 ± 1.60 | * | 34.57 ± 2.46 | *** |
| *swsn-9* | 16.25 ± 0.44 | ns | 16.46 ± 0.78 | ns | 16.00 ± 0.00 | ns | 15.96 ± 0.20 | ns | 32.42 ± 0.83 | ns | 32.25 ± 0.44 | ns |
| *swsn-2.1* | 16.33 ± 1.09 | ns | 18.05 ± 2.56 | ** | 16.27 ± 0.58 | ** | 17.14 ± 2.47 | * | 32.60 ± 1.38 | ns | 35.19 ± 3.76 | ** |
| *swsn-2.2* | 16.00 ± 0.00 | ns | 16.62 ± 0.74 | ns | 16.00 ± 0.00 | ns | 16.05 ± 0.22 | ns | 32.00 ± 0.00 | ns | 32.67 ± 0.80 | ns |
| *swsn-3* | 16.10 ± 0.31 | ns | 17.65 ± 1.63 | ** | 16.05 ± 0.22 | ns | 18.80 ± 3.72 | *** | 32.15 ± 0.49 | ns | 36.45 ± 4.06 | **** |
| *swsn-6* | 16.10 ± 0.64 | ns | 17.24 ± 1.41 | ns | 17.60 ± 3.15 | ** | 16.90 ± 1.70 | ** | 33.70 ± 3.16 | ** | 34.14 ± 2.18 | ** |
| *bcl-7* | 16.18 ± 0.39 | ns | 16.08 ± 0.40 | ns | 16.00 ± 0.00 | ns | 16.00 ± 0.00 | ns | 32.18 ± 0.39 | ns | 32.08 ± 0.40 | ns |
| *dpff-1* | 16.05 ± 0.22 | ns | 16.73 ± 0.88 | ns | 16.00 ± 0.00 | ns | 16.00 ± 0.00 | ns | 32.05 ± 0.22 | ns | 32.73 ± 0.88 | ns |
| *phf-10* | 16.16 ± 0.47 | ns | 18.52 ± 2.68 | ** | 16.00 ± 0.00 | ns | 17.78 ± 2.04 | **** | 32.16 ± 0.47 | ns | 36.30 ± 4.06 | **** |

**Supplementary Figure S1.** The SWI/SNF BAF complex promotes cell cycle exit. (**a**)The SWI/SNF complex consists of core, accessory, and BAF and PBAF-specific signature subunits. Table of *C. elegans* names used in this publication, alternative *C. elegans* nomenclature, and commonly used mammalian homologue names for the different subunits and (**b**) a schematic representation of the two conserved SWI/SNF subcomplexes (using *C. elegans* nomenclature). (**c**) Quantification of mesoblast lineage descendants, in the tail area (early dividing body wall muscles) and around the vulva (late dividing egg-laying muscles) at the L4 larval stage following RNAi by feeding of synchronised L1 larvae for the indicated genes, in wild type or *fzr-1* mutant backgrounds, with (**d**) table of mean mesoblast cell numbers, standard deviations and tests for significance compared to wild type and fzr-1 knockout larvae for each RNAi treatment. L4440 indicates the empty vector used in the RNAi control-treated animals.

Van der Vaart_Fig S2

**a**

```
C. elegans SWSN-4    ILADEMGLGKTIQTISLV
H. sapiens BRG1      ILADEMGLGKTIQTIALI
S. cerevisiae SWI2   ILADEMGLGKTIQTISLL
Drosophila brm       ILADEMGLGKTIQTISLV
```

**b**

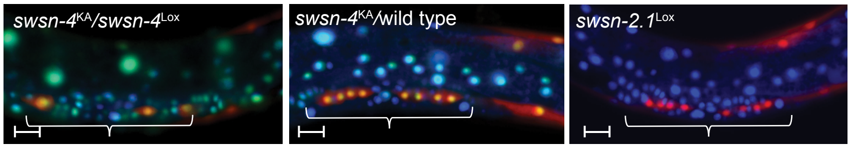


**c**

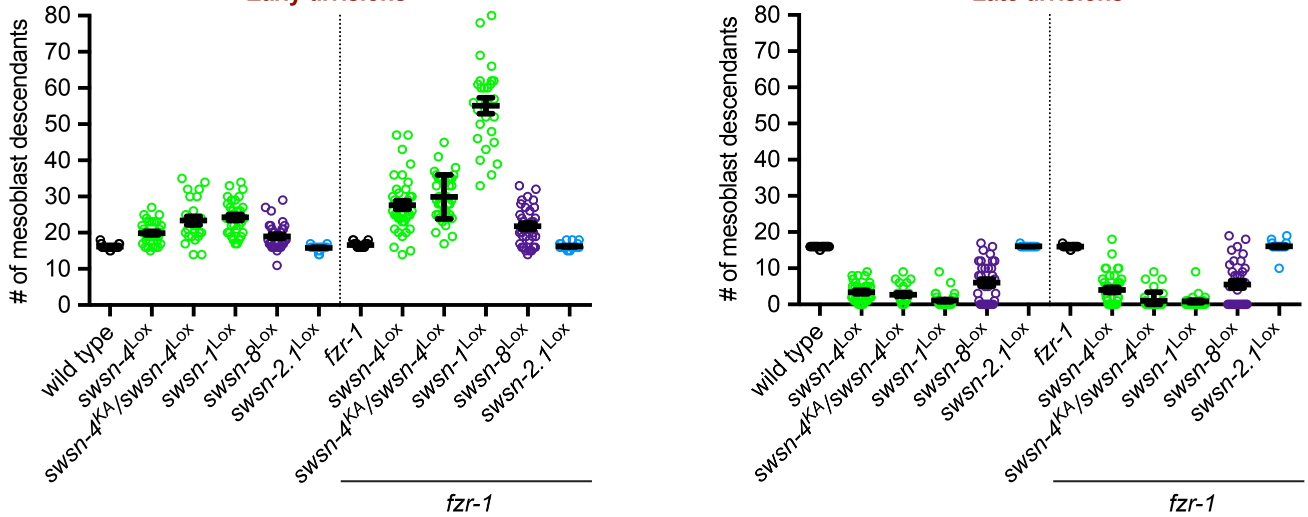


**d**

| Lox mutant | # cells around the vulva (late muscle) wild type | sig. | fzr-1 | sig. | # cells in the tail (early muscle) wild type | sig. | fzr-1 | sig. | total # of cells wild type | sig. | fzr-1 | sig. |
|---|---|---|---|---|---|---|---|---|---|---|---|---|
| wild type | 15.94 ± 0.25 | | 15.96 ± 0.35 | | 16.13 ± 0.56 | | 16.64 ± 0.70 | | 32.06 ± 0.63 | | 32.60 ± 0.76 | |
| *swsn-4* | 3.41 ± 2.72 | **** | 4.00 ± 4.43 | **** | 19.91 ± 3.01 | **** | 27.65 ± 7.58 | **** | 23.32 ± 3.87 | **** | 31.65 ± 7.32 | ns |
| *swsn-4^KA^ / swsn-4* Lox | 2.72 ± 2.56 | **** | 1.03 ± 2.36 | **** | 23.36 ± 5.92 | **** | 29.95 ± 6.08 | **** | 26.08 ± 5.22 | **** | 31.22 ± 6.28 | ns |
| *swsn-4^KA^* /wild type | 16.27 ± 1.57 | ns | 16.00 ± 0.00 | ns | 16.36 ± 1.45 | ns | 16.53 ± 0.68 | ns | 32.64 ± 2.98 | ns | 32.53 ± 0.68 | ns |
| *swsn-1* | 1.06 ± 1.97 | **** | 0.81 ± 1.82 | **** | 24.24 ± 4.68 | **** | 55.11 ± 11.63 | **** | 25.30 ± 5.00 | **** | 55.93 ± 11.19 | **** |
| *swsn-8* | 6.03 ± 5.57 | **** | 5.54 ± 6.12 | **** | 19.03 ± 3.58 | **** | 21.80 ± 5.28 | **** | 25.06 ± 5.41 | **** | 27.34 ± 6.61 | *** |
| *swsn-2.1* | 16.04 ± 0.19 | ns | 16.04 ± 1.43 | ns | 15.85 ± 0.66 | ns | 16.23 ± 0.82 | ns | 31.89 ± 0.70 | ns | 32.27 ± 1.82 | ns |

| |
|---|
| core |
| BAF |
| accessory |

**Supplementary Figure S2.** Simultaneous knockout of *fzr-1^Cdh1^* increases M descendant hyperplasia early after SWI/SNF gene knockout, but does not suppress the proliferation defect of late M descendants. (**a**) Illustration of the conserved lysine residue in the ATPase domain of SWSN-4, which is mutated to alanine (KA) to abolish ATPase activity. (**b**) Representative images of *swsn-4^KA^* trans-heterozygotes in the presence and absence of a wild type copy of *swsn-4* (marked in cis with P*eft-3::gfp*) and of *swsn-2.1^Lox^* knockout larvae. Brackets indicate egg-laying muscle precursors, scale bar 10 µm. (**c**) Quantification of mesoblast lineage descendants in the tail area (early formed BWM) and around the vulva (late dividing egg-laying muscle precursors) at the L4 larval stage of the indicated genotypes in wild type or *fzr-1* mutant backgrounds, with (**d**) table of mean mesoblast cell numbers, standard deviations and tests for significance compared to wild type and *fzr-1* knockout larvae for the indicated genotypes.

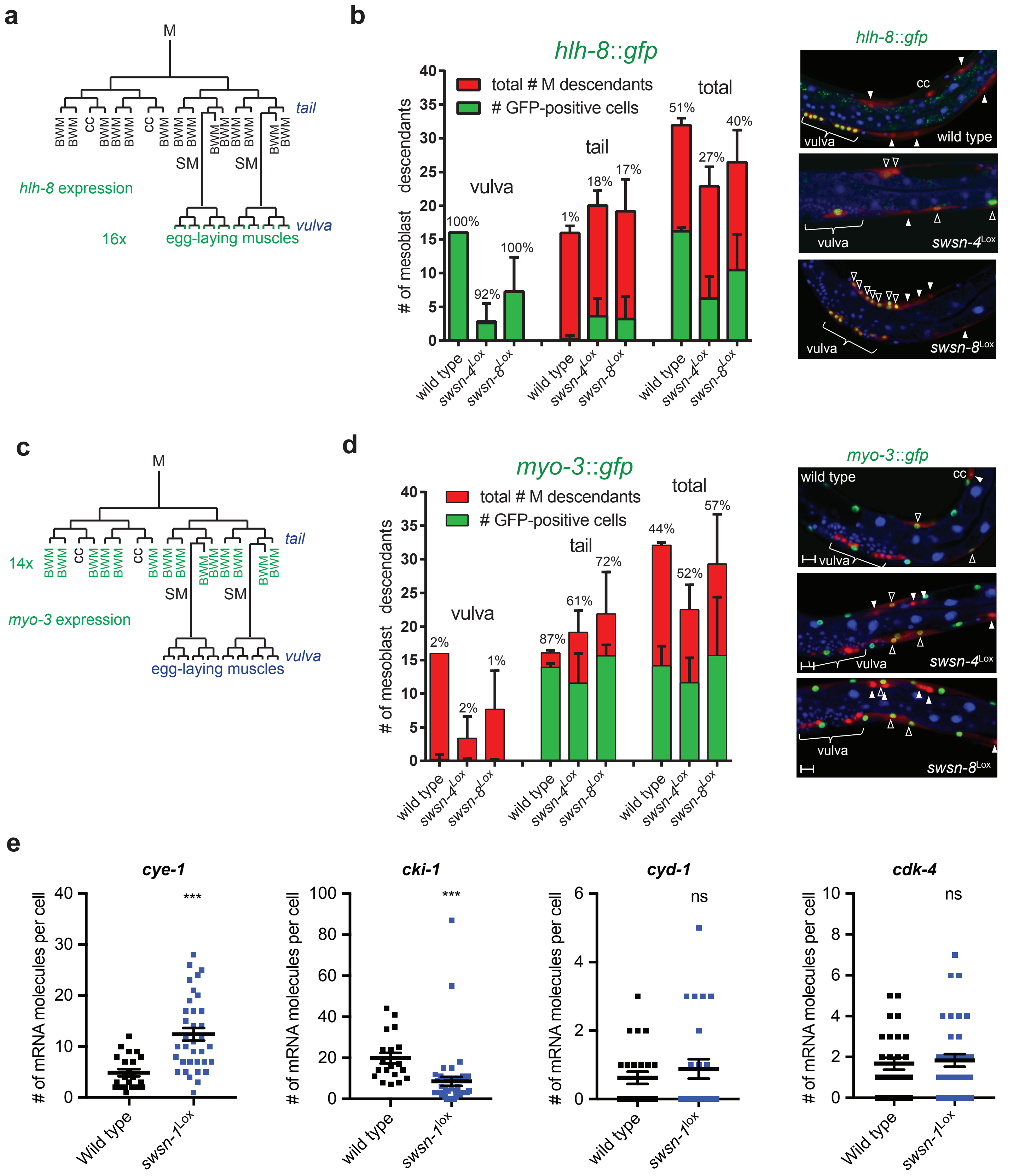


**Supplementary Figure S3.** Extra M lineage descendants in the conditional SWI/SNF knockout animals correspond to a prolonged proliferative undifferentiated state. (**a**) Illustration of the wild type expression pattern of P*hlh-8::gfp*, which is restricted to the 16 egg-laying muscle precursors in early L4 larvae. (**b**) Quantification of the number of mesoblast descendants expressing P*hlh-8::gfp* in early L4 larvae, for the indicated genotypes, with representative fluorescence microscopy images (right). Bars indicate the numbers of M descendants in the vulva region, tail, or total, with percentage of GFP-expressing cells indicated above the bars. Microscopy images: closed triangles indicate M descendants that only express the mCherry reporter. Open triangles indicate GFP expression in M descendants. Scale bar 10 µm. (**c**) Illustration of the wild type expression pattern of P*myo-3::gfp* in early L4 mesoblast descendants. (**d**) Quantification of number of mesoblast descendants expressing P*myo-3::gfp* in early L4 larvae, for the indicated genotypes, with representative images as under b. (**e**) Quantification of numbers of mRNA molecules per cell in smFISH experiments of early L4 wild-type and *swsn-1* knockout larvae for the indicated genes.

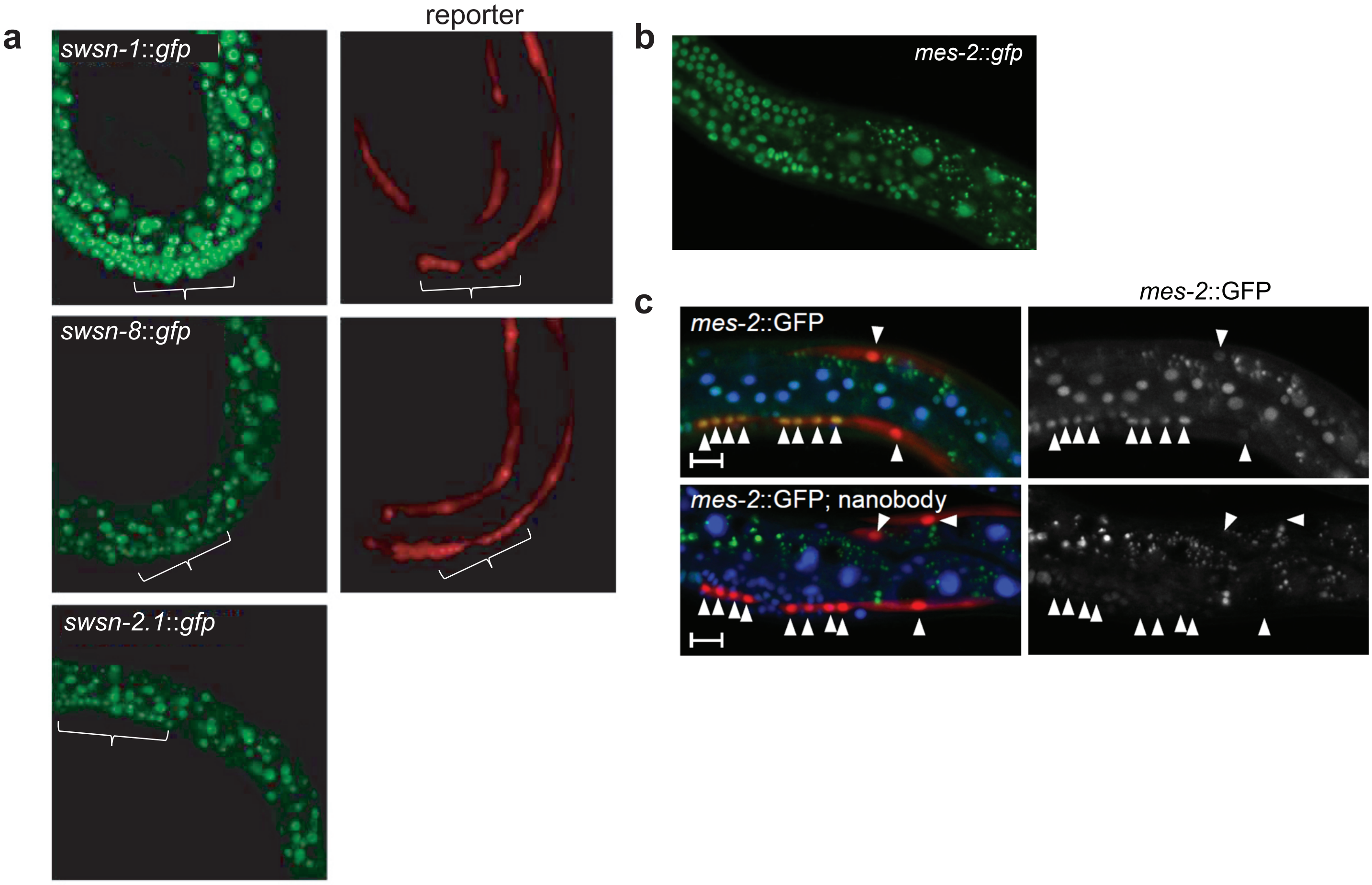


**Supplementary Figure S4.** Expression of endogenous GFP-tagged SWI/SNF subunits *swsn-1, swsn-8* and *swsn-2.1* as well as EZH2-related PcG gene *mes-2*. (**a**) Representative images of L4 larvae expressing endogenous GFP-tagged SWI/SNF subunits *swsn-1*, *swsn-8* and *swsn-2.1*. Brackets indicate late egg-laying muscle precursors. (**b**) Representative image of endogenous GFP-tagged EZH2-related PcG gene *mes-2*. (**c**) Expression of MES-2::GFP in L4 larvae in wild type worms (top) and after protein degradation (bottom, nanobody). Arrowheads indicate M lineage descendants.

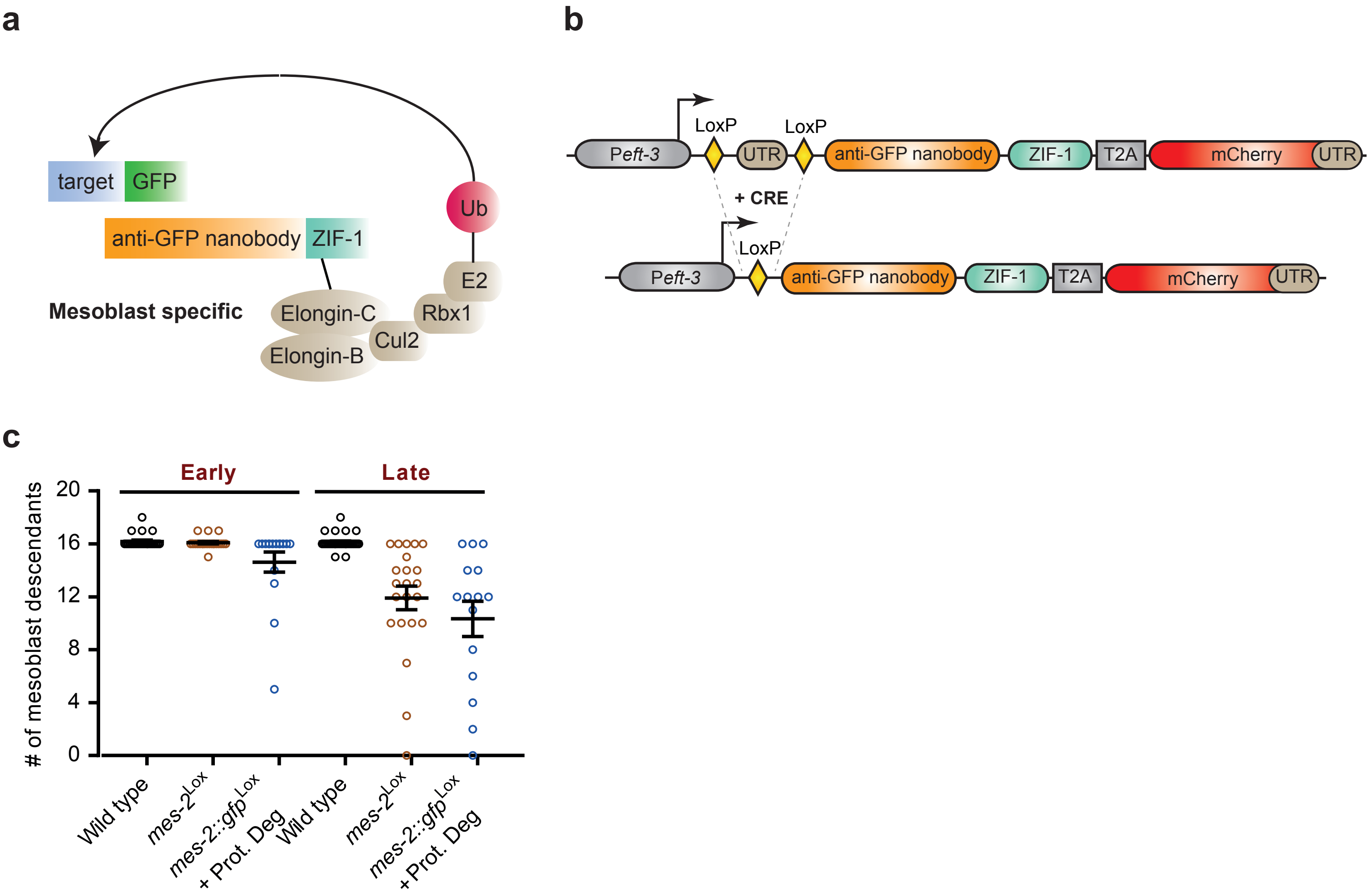


**Supplementary Figure S5.** Gene excision of *mes-2::gfp*$^{Lox}$ combined with protein degradation did not interfere with early M cell divisions. (**a**,**b**) Schematic of tissue-specific protein degradation strategy (**a**) an anti-GFP nanobody fused to the ZIF-1 subunit of a CUL-2 E3 ubiquitin ligase leads to specific degradation of GFP-tagged proteins (Prot. Deg.) (**b**) Schematic of the strategy for tissue-specific expression of the anti-GFP nanobody-ZIF-1 fusion protein. A floxed STOP cassette (LoxP::*let-858* UTR::LoxP) is inserted before the coding sequences of the fusion protein. Lineage-specific CRE recombinase expression leads to excision of the STOP cassette and nanobody::ZIF-1 expression from the general *eft-3* promoter. (**c**) Quantifications of mesoblast descendants for the indicated genotypes, in the tail area (early) and around the vulva (late).

**Supplementary Table 1**

| | |
|---|---|
| SV1863 | EG8082 heSi208 [Peft-3::LoxP::NLS egl-13::tagBFP2::tbb-2 UTR::LoxP::NLS egl-13::mCherry:: tbb-2 UTR]V; heSi141 [Phlh-8::CRE]X |
| SV1888 | EG8080 heSi210 [Peft-3::LoxP::NLS egl-13::tagBFP2::tbb-2 UTR::LoxP::NLS egl-13::mCherry:: tbb-2 UTR]III; (heSi141[Phlh-8::CRE]X |
| SV1930 | *swsn-8* (he273 [loxN exon3])I; *swsn-8* (he287 [loxN last intron])I; heSi208 [Peft-3::LoxP::NLS egl-13:: tagBFP2::tbb-2 UTR::LoxP::NLS egl-13::mCherry:: tbb-2 UTR]V; heSi141[Phlh-8::CRE]X; *pha-1* (+) III |
| SV1871 | *swsn-4* (he268 [LoxN start]); *swsn-4* (he272 [LoxN intron 5])IV; heSi208 [Peft-3::LoxP::NLS egl-13:: tagBFP2::tbb-2 UTR::LoxP::NLS egl-13::mCherry:: tbb-2 UTR]V; heSi141[Phlh-8::CRE]X; *pha-1* (+)III |
| SV1897 | *unc-119* (ed3)III; oxTi970 [eft-3p::GFP::2xNLS::tbb-2 3'UTR + Cbr-unc-119(+)]IV; *swsn-4* (he277[K564A]); *swsn-4* (he268 [LoxN start]; *swsn-4* (he272 [LoxN intron 5])IV; heSi208 [Peft-3::LoxP::NLS egl-13::tagBFP2::tbb-2 UTR::LoxP::NLS egl-13:: mCherry:: tbb-2 UTR]V; heSi141[Phlh-8::CRE]X; *pha-1* (+)III |
| SV2044 | *unc-119* (ed3)III; oxTi970 [eft-3p::GFP::2xNLS::tbb-2 3'UTR + Cbr-unc-119(+)]IV; *swsn-4* (he277[K564A]); *swsn-4* (+)IV; heSi208 [Peft-3::LoxP::NLS egl-13::tagBFP2::tbb-2 UTR::LoxP::NLS egl-13:: mCherry:: tbb-2 UTR]V; heSi141[Phlh-8::CRE]X |
| SV2025 | *swsn-2.1* (he309[swsn-2.1 LoxN first intron + LoxN last intron]) III; heSi210 [Peft-3::LoxP::NLS egl-13::tagBFP2::tbb-2 UTR::LoxP::NLS egl-13:: mCherry:: tbb-2 UTR]III; heSi208 [Peft-3::LoxP::NLS egl-13::tagBFP2::tbb-2 UTR::LoxP::NLS egl-13:: mCherry:: tbb-2 UTR]V; heSi141[Phlh-8::CRE])X *pha-1* (+)III |
| SV1941 | *fzr-1* (ku298)II; *unc-4* (e120)II; *swsn-8* (he273 [loxN exon3]) I; *swsn-8* ( (he287 [loxN last intron])I; heSi208 [Peft-3::LoxP::NLS egl-13:: tagBFP2::tbb-2 UTR::LoxP::NLS egl-13::mCherry:: tbb-2 UTR]V; heSi141[Phlh-8::CRE]X; *pha-1* (+)III |
| SV1894 | *fzr-1* (ku298)II; *unc-4* (e120)II; *swsn-4* (he268 [LoxN start]; *swsn-4* he272 [LoxN intron 5])IV; heSi208 [Peft-3::LoxP::NLS egl-13:: tagBFP2::tbb-2 UTR::LoxP::NLS egl-13::mCherry:: tbb-2 UTR]V; heSi141[Phlh-8::CRE]X; *pha-1* (+)III |
| SV2034 | *fzr-1* (ku298)II; *unc-4* (e120)II; *unc-119* (ed3)III; oxTi970 [eft-3p::GFP::2xNLS::tbb-2 3'UTR + Cbr-unc-119(+)]IV; *swsn-4* (he277[K564A]); *swsn-4* (he268 [LoxN start]; *swsn-4* (he272 [LoxN intron 5])IV; heSi208 [Peft-3::LoxP::NLS egl-13::tagBFP2::tbb-2 UTR::LoxP::NLS egl-13:: mCherry:: tbb-2 UTR]V; heSi141[Phlh-8::CRE]X; *pha-1* (+)III |
| SV2033 | *fzr-1* (ku298)II; *unc-4* (e120)II; *unc-119* (ed3)III; oxTi970 [eft-3p::GFP::2xNLS::tbb-2 3'UTR + Cbr-unc-119(+)]IV; *swsn-4* (he277[K564A]); *swsn-4* (+)IV; heSi208 [Peft-3::LoxP::NLS egl-13::tagBFP2::tbb-2 UTR::LoxP::NLS egl-13:: mCherry:: tbb-2 UTR]V; heSi141[Phlh-8::CRE]X |
| SV2029 | *fzr-1* (ku298)II; *unc-4* (e120)II; *swsn-2.1* (he309[swsn-2.1 LoxN first intron + LoxN last intron]) III; heSi210 [Peft-3::LoxP::NLS egl-13::tagBFP2::tbb-2 UTR::LoxP::NLS egl-13::mCherry:: tbb-2 UTR]III; heSi141[Phlh-8::CRE] X; pha-1(+)III |
| SV2027 | EG8080 heSi219 [Phlh-8::GFP::H2B::unc-54 UTR]III; heSi208 [Peft-3::LoxP::NLS egl-13::tagBFP2::tbb-2 UTR::LoxP::NLS egl-13::mCherry:: tbb-2 UTR]V; heSi141 [hlh-8::CRE]X |
| SV2030 | *swsn-4* (he268 [LoxN start]); *swsn-4* (he272 [LoxN intron 5])IV; heSi219 [Phlh-8::GFP::H2B::unc-54 UTR]III; heSi208 [Peft-3::LoxP::NLS egl-13::tagBFP2::tbb-2 UTR::LoxP::NLS egl-13::mCherry:: tbb-2 UTR]V; heSi141 [hlh-8::CRE]X |

| | |
|---|---|
| SV2031 | *swsn-8* (he273 [loxN exon3])I; *swsn-8* (he287 [loxN last intron])I; heSi219 [Phlh-8::GFP::H2B::unc-54 UTR]III; heSi208 [Peft-3::LoxP::NLS egl-13::tagBFP2::tbb-2 UTR::LoxP::NLS egl-13::mCherry:: tbb-2 UTR]V; heSi141 [hlh-8::CRE]X |
| SV1578 | unc-119(ed3)III; heIs105 [rps-27::loxP::nls::mCherry::let-858UTR::loxP::nls::gfp::let-858 UTR]IV; swsn-1(os22)V; heSi164 [rps-27::loxN::nls::mCherry::let-858 UTR::Pswsn-1::swsn-1::unc-54 UTR::loxN::nls::gfp::let-858UTR]V; heSi141 [hlh-8::CRE]X |
| SV2032 | unc-119(ed3)III; heIs105 [rps-27::loxP::nls::mCherry::let-858UTR::loxP::nls::gfp::let-858 UTR]IV; swsn-1(os22)V; heSi164 [rps-27::loxN::nls::mCherry::let-858 UTR::Pswsn-1::swsn-1::unc-54 UTR::loxN::nls::gfp::let-858UTR]V; heSi141 [hlh-8::CRE]X; heEx611 [hlh-8::hairpin swsn-1::unc-54] |
| SV1818 | pha-1(e2123)III 7x out-crossed |
| SV2023 | swsn-1 (he306[swsn-1::eGFP])V; heSi208[blue to red read-out]V; heSi141[hlh-8::CRE]X |
| SV2077 | swsn-1 (he313[swsn-1::eGFP::LoxP])V; swsn-1 (he315[LoxP second intron])V; heSi208 [Peft-3::LoxP::NLS egl-13::tagBFP2::3'UTR tbb-2::LoxP::NLS egl-13::mCherry::3'UTR tbb-2] V; (heSi141[Phlh-8::CRE])X |
| SV2098 | fzr-1(ku298;unc-4(e120)II; swsn-1 (he313[swsn-1::eGFP::LoxP])V; swsn-1 (he315[LoxP second intron])V; heSi208 [Peft-3::LoxP::NLS egl-13::tagBFP2::3'UTR tbb-2::LoxP::NLS egl-13::mCherry::3'UTR tbb-2] V; (heSi141[Phlh-8::CRE])X |
| SV2067 | heSi210 [Peft-3::LoxP::NLS egl-13::tagBFP2::tbb-2 UTR::LoxP::NLS egl-13::mCherry:: tbb-2 UTR]III; (heSi141[Phlh-8::CRE]X; he303[peft-3-LoxP-let-858 UTR-LoxP vhh gfp::ZIF-1mCherry]) IV ; swsn-1 (he313[swsn-1::GFP::LoxP] ; swsn-1 (he315[LoxP second intron])V ; heSi141 [hlh-8::cre] X |
| SV2161 | heSi210 [Peft-3::LoxP::NLS egl-13::tagBFP2::tbb-2 UTR::LoxP::NLS egl-13::mCherry:: tbb-2 UTR]III; swsn-1 (he313[swsn-1::eGFP::LoxP])V; heSi208 [Peft-3::LoxP::NLS egl-13::tagBFP2::3'UTR tbb-2::LoxP::NLS egl-13::mCherry::3'UTR tbb-2] V; (heSi141[Phlh-8::CRE])X |
| SV2155 | mes-2 (he332[loxN first intron]); mes-2 (he340[loxN last intron]) II; heSi208 [Peft-3::LoxP::NLS egl-13::tagBFP2::3'UTR tbb-2::LoxP::NLS egl-13::mCherry::3'UTR tbb-2] V; (heSi141[Phlh-8::CRE])X |
| SV2185 | mes-2 (he332[loxN first intron]); mes-2 (he340[loxN last intron]) II; *swsn-4* (he268 [LoxN start]); *swsn-4* (he272 [LoxN intron 5])IV; heSi208 [Peft-3::LoxP::NLS egl-13::tagBFP2::3'UTR tbb-2::LoxP::NLS egl-13::mCherry::3'UTR tbb-2] V; (heSi141[Phlh-8::CRE])X |
| SV2186 | *swsn-8* (he273 [loxN exon3])I; *swsn-8* (he287 [loxN last intron])I; mes-2 (he332[loxN first intron]); mes-2 (he340[loxN last intron]) II; heSi208 [Peft-3::LoxP::NLS egl-13::tagBFP2::3'UTR tbb-2::LoxP::NLS egl-13::mCherry::3'UTR tbb-2] V; (heSi141[Phlh-8::CRE])X |
| SV2187 | mes-2 (he332[loxN first intron]); mes-2 (he340[loxN last intron]) II; swsn-1 (he313[swsn-1::eGFP::LoxP])V; swsn-1 (he315[LoxP second intron]) V; heSi208 [Peft-3::LoxP::NLS egl-13::tagBFP2::3'UTR tbb-2::LoxP::NLS egl-13::mCherry::3'UTR tbb-2] V; (heSi141[Phlh-8::CRE])X |
| SV2105 | cki-1(he329[loxP first intron, loxP in UTR]) II, heSi208 [Peft-3::LoxP::NLS egl-13::tagBFP2::3'UTR tbb-2::LoxP::NLS egl-13::mCherry::3'UTR tbb-2] V; (heSi141[Phlh-8::CRE]); ayIS6[Phlh-8::GFP])X |
| SV2122 | cki-1(he329[loxP first intron, loxP in UTR]) II, *swsn-4* (he277[K564A]); *swsn-4* (+)IV heSi208 [Peft-3::LoxP::NLS egl-13::tagBFP2::3'UTR tbb-2::LoxP::NLS egl-13::mCherry::3'UTR tbb-2] V; (heSi141[Phlh-8::CRE]); ayIS6[Phlh-8::GFP])X |

| | |
|---|---|
| SV2195 | mes-2 (he332[loxN first intron]); mes-2 (he342[mes-2::eGFP::LoxN]) II, heSi210 [Peft-3::LoxP::NLS egl-13::tagBFP2::tbb-2 UTR::LoxP::NLS egl-13::mCherry:: tbb-2 UTR]III; (heSi141[Phlh-8::CRE]X; he303[peft-3-LoxP-let-858 UTR-LoxP vhh gfp::ZIF-1mCherry]) IV ; heSi141 [hlh-8::cre] X |
| SV2196 | mes-2 (he332[loxN first intron]); mes-2 (he342[mes-2::eGFP::LoxN]) II, heSi210 [Peft-3::LoxP::NLS egl-13::tagBFP2::tbb-2 UTR::LoxP::NLS egl-13::mCherry:: tbb-2 UTR]III; (heSi141[Phlh-8::CRE]X; he303[peft-3-LoxP-let-858 UTR-LoxP vhh gfp::ZIF-1mCherry]) IV ; swsn-1 (he313[swsn-1::eGFP::LoxP] ; swsn-1 (he315[LoxP second intron]) V ; heSi141 [hlh-8::cre] X |

**Supplementary Table 2**

| Wormbase sequence name | baseMean | log2FoldChange | lfcSE | stat |
|---|---|---|---|---|
| F59D12.2 | 3,553001723 | -5,19639252 | 1,113035484 | 4,668667435 |
| R05C11.1 | 2,223615823 | -4,562274215 | 1,199823217 | 3,802455355 |
| C28F5.2 | 6,429409582 | -4,511729685 | 0,913031585 | 4,941482595 |
| F32D8.10 | 3,228746951 | -4,470295084 | 1,274441407 | 3,507650536 |
| F43C9.2 | 7,434220571 | -4,388287121 | 0,968117557 | 4,532803983 |
| F10E9.12 | 1,933005356 | -4,299582936 | 1,217128709 | 3,532562253 |
| ZK525.1 | 164,9808284 | -4,19806805 | 0,359195269 | 11,68742578 |
| F31D4.8 | 16,88529544 | -4,142960775 | 0,590332424 | 7,018013255 |
| C52B9.1 | 6,275960445 | -4,001446199 | 0,900091509 | 4,445599319 |
| T25F10.3 | 2,406403021 | -3,965704097 | 1,157928634 | 3,424826005 |
| C50D2.2 | 2,324716666 | -3,938937314 | 1,112386651 | 3,540978589 |
| T01B10.4 | 2,042184 | -3,8347596 | 1,2839331 | 2,986729 |
| F16B4.8 | 17,00525359 | -3,823443111 | 0,592774596 | 6,450079233 |
| F35H10.10 | 6,826735104 | -3,744602494 | 0,800837673 | 4,675857071 |
| T28F2.3 | 2,91380979 | -3,723118201 | 1,129486406 | 3,296293059 |
| T01D3.7 | 4,931104361 | -3,701223338 | 1,052948671 | 3,515103292 |
| F55C12.4 | 2,508801674 | -3,575991038 | 1,067259634 | 3,350628961 |
| T23G11.6 | 2,263758 | -3,4926988 | 1,1143712 | 3,134233 |
| C49F8.3 | 1,834561 | -3,4892562 | 1,1801507 | 2,956619 |
| Y73B6BL.9 | 117,6541599 | -3,473782358 | 0,331770405 | 10,47044072 |
| ZK337.2 | 7,51285056 | -3,403759518 | 0,708544461 | 4,803875702 |
| F53C3.12 | 2,30234 | -3,342929 | 1,1558272 | 2,892239 |
| Y51A2D.15 | 3,717869271 | -3,321991113 | 0,930150795 | 3,57145436 |
| C30F12.1 | 5,052848278 | -3,280743066 | 0,847869627 | 3,869395675 |
| W10G6.3 | 2,810952 | -3,251047 | 1,0950925 | 2,968742 |
| Y75B7AL.1 | 5,358564833 | -3,235368961 | 0,846432309 | 3,822359951 |
| T05A8.3 | 17,20491782 | -3,233004597 | 0,876275617 | 3,689483688 |
| C53C11.3 | 3,536610706 | -3,209639388 | 1,001886115 | 3,203597034 |
| R05H11.2 | 80,33633224 | -3,081906221 | 0,264943538 | 11,63231321 |
| C06H2.5 | 17,89604657 | -3,004433597 | 0,473712632 | 6,342312599 |
| C18A3.4 | 13,33058858 | -2,970176766 | 0,695218308 | 4,272293658 |
| T09B4.5 | 38,69459771 | -2,923290406 | 0,489642049 | 5,970260131 |
| C05G5.7 | 20,24214652 | -2,902575221 | 0,491125568 | 5,910047068 |
| T06D8.1 | 4,799309114 | -2,886163471 | 0,871758441 | 3,310737624 |
| T04F8.9 | 5,512121889 | -2,864457451 | 0,777926522 | 3,682169677 |
| F15G9.5 | 7,274285883 | -2,846746795 | 0,656899 | 4,33361414 |
| C30G7.2 | 5,187067201 | -2,843805398 | 0,849502784 | 3,347611627 |
| T02C12.1 | 8,18477688 | -2,833099859 | 0,662045696 | 4,279311646 |
| T21D12.9 | 5,534132705 | -2,80116694 | 0,737610555 | 3,797623175 |
| T05A8.8 | 5,121132937 | -2,789422465 | 0,708652526 | 3,936234422 |
| F32H2.5 | 7,452785005 | -2,77429848 | 0,66735277 | 4,157169348 |
| Y39A3CL.2 | 4,587384 | -2,7483961 | 0,8946785 | 3,071937 |
| K07D4.8 | 13,37302059 | -2,724567812 | 0,508755069 | 5,355362484 |
| T19D12.6 | 4,1306 | -2,6874172 | 0,9202694 | 2,92025 |
| Y22D7AL.8 | 3,460784 | -2,6471055 | 0,8969722 | 2,951157 |
| F19B10.10 | 21,37065711 | -2,642869321 | 0,390570219 | 6,766694415 |
| ZK909.4 | 3,411797 | -2,6322978 | 0,8645328 | 3,044763 |
| C54G4.5 | 3,21844 | -2,6223438 | 0,8694378 | 3,016137 |

| | | | | |
|---|---|---|---|---|
| F49E10.5 | 3,568208 | -2,5731958 | 0,8399179 | 3,063628 |
| C32D5.7 | 4,455509724 | -2,553599262 | 0,775687108 | 3,292048088 |
| C44F1.3 | 28,00099766 | -2,525912915 | 0,374841678 | 6,738612758 |
| Y71G12B.16 | 15,13765716 | -2,470299758 | 0,431020702 | 5,731278678 |
| C04E12.7 | 5,044876013 | -2,348057057 | 0,655091383 | 3,584319863 |
| F45E1.2 | 4,844730404 | -2,322006929 | 0,709461605 | 3,272914154 |
| D2092.6 | 7,659303957 | -2,305441036 | 0,59816141 | 3,854212252 |
| F11C1.6 | 6,53651 | -2,30243 | 0,7376172 | 3,121443 |
| F26D11.10 | 4,862096364 | -2,270404165 | 0,687909503 | 3,30044018 |
| W08A12.3 | 5,678335066 | -2,261343597 | 0,675965833 | 3,345351922 |
| F25H5.1 | 15,11377094 | -2,241816015 | 0,631308261 | 3,551063961 |
| C39E6.4 | 155,9913717 | -2,235106212 | 0,239336654 | 9,338754335 |
| M05B5.2 | 9,001235006 | -2,21446837 | 0,513202536 | 4,314998884 |
| C44C11.1 | 91,14952856 | -2,204667552 | 0,254394587 | 8,666330426 |
| R08C7.12 | 11,19121296 | -2,199944624 | 0,589763606 | 3,73021428 |
| T01E8.3 | 5,830174242 | -2,195306919 | 0,589544727 | 3,7237326 |
| F55C5.3 | 4,250925 | -2,1756469 | 0,747322 | 2,911257 |
| T19E7.6 | 4,445232 | -2,1736753 | 0,7269539 | 2,990115 |
| F08G2.7 | 7,011230173 | -2,166141491 | 0,648571461 | 3,33986557 |
| R02E12.5 | 7,91289554 | -2,122702915 | 0,59226882 | 3,584019358 |
| ZC132.4 | 20,3037772 | -2,100639436 | 0,504947468 | 4,160114802 |
| F49E12.6 | 35,85041221 | -2,082593893 | 0,308107176 | 6,759316417 |
| H27C11.1 | 10,75765713 | -2,070573489 | 0,598890432 | 3,457349422 |
| Y65B4BL.1 | 9,478951458 | -2,049321124 | 0,500525304 | 4,0943407 |
| F46C8.7 | 19,04005952 | -2,04531589 | 0,50710685 | 4,03330361 |
| Y38C1BA.2 | 4,930628 | -2,0404611 | 0,642023 | 3,178174 |
| C50C3.9 | 6,334574816 | -2,028039188 | 0,577940788 | 3,509077797 |
| C34G6.4 | 6,206016 | -2,0231026 | 0,6562789 | 3,082687 |
| F19B2.5 | 7,291303 | -2,0210459 | 0,6850437 | 2,950244 |
| F28B4.2 | 10,72974931 | -2,001002284 | 0,467929135 | 4,276293423 |
| C46H11.11 | 11,98527244 | -1,989432696 | 0,485534026 | 4,097411488 |
| K12F2.2 | 12,49982927 | -1,986023233 | 0,416800123 | 4,764929569 |
| T22C8.8 | 7,298187373 | -1,969957582 | 0,566171988 | 3,479433145 |
| F55A4.2 | 12,98813294 | -1,960766987 | 0,484358053 | 4,048176707 |
| T13H5.6 | 5,009567 | -1,9595866 | 0,6775986 | 2,891958 |
| F44G4.8 | 8,501526 | -1,9427094 | 0,6242212 | 3,112213 |
| F21D5.9 | 14,25814149 | -1,940913328 | 0,470391294 | 4,126167622 |
| F58H10.1 | 11,738375 | -1,9400965 | 0,6422476 | 3,020792 |
| F22F1.1 | 277,3998071 | -1,92113174 | 0,31168151 | 6,163765505 |
| F44E2.10 | 6,213664625 | -1,84274245 | 0,560391001 | 3,288315566 |
| K08E5.2 | 10,06448743 | -1,835986393 | 0,507869617 | 3,615074289 |
| F08C6.7 | 19,21326545 | -1,830291839 | 0,339714511 | 5,387735229 |
| T10B10.4 | 78,86275841 | -1,806117404 | 0,222848952 | 8,104670853 |
| C13C4.4 | 6,77314 | -1,7916252 | 0,5638847 | 3,17729 |
| F07C6.4 | 75,83796827 | -1,790007424 | 0,287191287 | 6,232805474 |
| F42G4.3 | 27,3709441 | -1,789843729 | 0,451434187 | 3,964794383 |
| F37A8.5 | 76,23856424 | -1,788332121 | 0,427076354 | 4,187382661 |
| W02D9.10 | 231,826969 | -1,780775628 | 0,174146013 | 10,22576169 |
| W02B8.1 | 11,14598895 | -1,739374501 | 0,45727691 | 3,803766304 |
| F33C8.3 | 13,39255702 | -1,731125269 | 0,442112941 | 3,915572487 |

| | | | | |
|---|---|---|---|---|
| R12H7.1 | 11,30291773 | -1,72862014 | 0,459953499 | 3,75824979 |
| F46H6.1 | 12,16983613 | -1,72561875 | 0,404885836 | 4,261988436 |
| F01G12.5 | 21,30316162 | -1,723359598 | 0,500334447 | 3,444415251 |
| K05G3.3 | 8,773102472 | -1,703444845 | 0,51339676 | 3,317989083 |
| F45E1.7 | 12,54623911 | -1,701288838 | 0,45570649 | 3,733299556 |
| Y54E5B.1 | 22,918993 | -1,70001222 | 0,378673612 | 4,489386555 |
| C48A7.1 | 20,4225952 | -1,697401474 | 0,480083201 | 3,53564022 |
| F26F4.5 | 69,362556 | -1,6942338 | 0,5324367 | 3,182038 |
| C33B4.3 | 20,15406795 | -1,677720195 | 0,447757266 | 3,74694131 |
| T12A7.2 | 14,94638619 | -1,674499295 | 0,4188446 | 3,997901123 |
| T22C1.7 | 16,03176992 | -1,670113273 | 0,518549407 | 3,220740877 |
| Y43F8B.2 | 131,6747732 | -1,666117167 | 0,297075851 | 5,608389775 |
| T06C10.4 | 14,93295356 | -1,658161752 | 0,395250783 | 4,195214336 |
| F25H8.3 | 68,0917646 | -1,645229059 | 0,369805425 | 4,448904609 |
| F42G8.11 | 7,494057 | -1,570831 | 0,5154163 | 3,047694 |
| Y15E3A.5 | 34,31500713 | -1,555501294 | 0,341134451 | 4,559789521 |
| T11F9.11 | 14,01838371 | -1,540876567 | 0,465300004 | 3,311576521 |
| F53G12.5 | 64,30633456 | -1,537568106 | 0,352784713 | 4,358375088 |
| C26E6.2 | 32,98989403 | -1,517440675 | 0,320675058 | 4,732019653 |
| C01F6.6 | 14,90318401 | -1,508234219 | 0,393978252 | 3,828216944 |
| C18H9.7 | 28,88622929 | -1,50694425 | 0,269820686 | 5,58498414 |
| Y71H10B.1 | 82,20567884 | -1,493053011 | 0,334182765 | 4,467773832 |
| T23G4.1 | 29,87975777 | -1,478702384 | 0,369616516 | 4,00063937 |
| C01C10.3 | 40,80348001 | -1,47086494 | 0,335903364 | 4,378833606 |
| B0034.3 | 22,668541 | -1,4276013 | 0,4612311 | 3,095198 |
| F54G8.3 | 21,53793518 | -1,426167136 | 0,367907844 | 3,876424925 |
| F14B4.2 | 47,19067659 | -1,419814141 | 0,304926294 | 4,656253563 |
| B0464.4 | 22,34895784 | -1,417670336 | 0,317629508 | 4,463282847 |
| C02B10.5 | 76,62444382 | -1,405198671 | 0,280465481 | 5,010237497 |
| F52D2.6 | 51,65223978 | -1,401390468 | 0,270383975 | 5,182964215 |
| Y22F5A.3 | 29,09373674 | -1,396804685 | 0,301955119 | 4,625868542 |
| T19H5.4 | 24,186515 | -1,3666714 | 0,4702452 | 2,906296 |
| T07A9.7 | 69,76875605 | -1,34177833 | 0,199796108 | 6,715738062 |
| ZK617.1 | 11,843322 | -1,3393497 | 0,4396667 | 3,046284 |
| C12C8.1 | 21,208646 | -1,3382157 | 0,4553537 | 2,938849 |
| T13B5.1 | 173,4074229 | -1,313326399 | 0,208300187 | 6,304969846 |
| Y47D3A.17 | 19,7880015 | -1,309650899 | 0,373042518 | 3,510728234 |
| W03G9.1 | 36,64803704 | -1,298534509 | 0,344545864 | 3,768829191 |
| R02E12.2 | 51,607165 | -1,2969935 | 0,4192553 | 3,093565 |
| H24G06.1 | 55,87921854 | -1,284911929 | 0,375855815 | 3,418629903 |
| C33A11.1 | 57,38034944 | -1,279591709 | 0,298301575 | 4,289590854 |
| T05H10.4 | 26,231066 | -1,276351535 | 0,321241824 | 3,973179829 |
| Y41C4A.13 | 57,53268353 | -1,274968515 | 0,212471588 | 6,000654151 |
| C17C3.1 | 23,71596561 | -1,274503774 | 0,349029798 | 3,651561498 |
| T06E6.2 | 289,64723 | -1,2610586 | 0,4138514 | 3,047129 |
| ZK637.1 | 42,23495338 | -1,230849931 | 0,355300656 | 3,464248973 |
| R03E9.3 | 38,92558198 | -1,230776863 | 0,279170951 | 4,408685287 |
| C45G3.1 | 25,87825617 | -1,220832368 | 0,282780522 | 4,317243487 |
| ZK524.2 | 50,93734804 | -1,204126006 | 0,362101484 | 3,325382687 |
| W09D6.5 | 86,33110252 | -1,18387838 | 0,196959176 | 6,010780538 |

| | | | | |
|---|---|---|---|---|
| F57F5.5 | 56,73284724 | -1,181781983 | 0,301234091 | 3,92313493 |
| B0365.3 | 121,5912869 | -1,172017285 | 0,354975512 | 3,301684895 |
| T14F9.4 | 49,05795247 | -1,171426729 | 0,299981099 | 3,905001784 |
| K08H10.2 | 289,3681352 | -1,170823142 | 0,269509268 | 4,344277851 |
| C06G3.2 | 30,91447711 | -1,164211552 | 0,307233604 | 3,789336636 |
| Y47D3A.6 | 19,15408759 | -1,155002958 | 0,345239775 | 3,34550953 |
| C53C11.5 | 14,167833 | -1,1482234 | 0,395382 | 2,904086 |
| Y56A3A.29 | 38,77535068 | -1,138770617 | 0,328704298 | 3,464422651 |
| F47F6.1 | 234,291006 | -1,127608941 | 0,352145876 | 3,20210747 |
| Y47D9A.2 | 34,982855 | -1,09523 | 0,3743581 | 2,925621 |
| T27F2.2 | 21,502562 | -1,0913504 | 0,3736086 | 2,921106 |
| C07A9.12 | 13,112497 | -1,0893231 | 0,3592731 | 3,03202 |
| T20D4.6 | 36,1342034 | -1,074502397 | 0,255816767 | 4,200281362 |
| W06F12.1 | 101,6322586 | -1,057639805 | 0,233696956 | 4,525689261 |
| AC7.2 | 25,54251865 | -1,037070247 | 0,2924635 | 3,545981797 |
| F55F1.1 | 76,51257812 | -1,037028655 | 0,197330676 | 5,255283561 |
| Y105C5B.21 | 36,63704 | -1,0324217 | 0,3565687 | 2,895435 |
| Y53G8AM.8 | 22,655643 | -1,0292847 | 0,3288706 | 3,129756 |
| F08B6.4 | 32,64467199 | -1,025459056 | 0,307366528 | 3,336274328 |
| C14B4.2 | 122,3799226 | -1,021443598 | 0,25068799 | 4,07456136 |
| F42G9.9 | 19,352785 | -1,019695 | 0,3294334 | 3,0953 |
| F08F1.8 | 22,177381 | -1,0171508 | 0,3298314 | 3,083851 |
| ZK381.5 | 54,38917595 | -1,007983872 | 0,217284179 | 4,639011814 |
| B0336.1 | 45,914659 | -0,9816407 | 0,3362004 | 2,919808 |
| C43E11.6 | 45,40564056 | -0,942323567 | 0,286001609 | 3,294819107 |
| W04D2.1 | 56,44284278 | -0,939779243 | 0,216087554 | 4,349066956 |
| C10E2.6 | 78,09540205 | -0,938889985 | 0,266879458 | 3,518030165 |
| C35D10.13 | 223,1614331 | -0,919307202 | 0,264842145 | 3,471151469 |
| K04H4.1 | 172,0875432 | -0,914601 | 0,252152868 | 3,627168732 |
| F52G3.1 | 79,93252937 | -0,903524703 | 0,265615381 | 3,401627949 |
| ZK177.6 | 97,57609651 | -0,902340595 | 0,200139992 | 4,508547164 |
| Y49E10.6 | 879,3923845 | -0,901202441 | 0,242029551 | 3,723522346 |
| ZK1151.1 | 213,511523 | -0,8990057 | 0,3101886 | 2,898256 |
| Y113G7B.23 | 422,381722 | -0,8882372 | 0,2805206 | 3,166389 |
| F43G9.9 | 21,338275 | -0,883303 | 0,2957611 | 2,986542 |
| F54B11.3 | 72,32708707 | -0,867567879 | 0,22002523 | 3,94303817 |
| Y59H11AR.2 | 104,3671546 | -0,854683371 | 0,248268241 | 3,442580359 |
| Y69A2AR.7 | 39,51510103 | -0,84977401 | 0,258845965 | 3,282933198 |
| F26B1.3 | 328,5412955 | -0,838990925 | 0,244902898 | 3,425810517 |
| F36D4.3 | 121,2538489 | -0,837784648 | 0,230257891 | 3,638462265 |
| F55G1.4 | 44,13139261 | -0,835766161 | 0,244865555 | 3,41316344 |
| F23B12.8 | 27,600287 | -0,8280302 | 0,2677484 | 3,092568 |
| T05G5.3 | 276,9496133 | -0,812336534 | 0,18823952 | 4,315440957 |
| ZK484.4 | 56,76769438 | -0,800572652 | 0,243514477 | 3,287577243 |
| C04C11.2 | 55,18676228 | -0,783962788 | 0,205143525 | 3,821533172 |
| F32A7.5 | 104,8367477 | -0,744738436 | 0,200945061 | 3,706179347 |
| Y69A2AR.30 | 328,0470597 | -0,742827956 | 0,19888859 | 3,734894766 |
| C26C6.5 | 79,89454876 | -0,732542857 | 0,204563252 | 3,581009045 |
| F28C1.3 | 45,26705959 | -0,728115587 | 0,226181132 | 3,219170319 |
| T09B9.4 | 50,86168534 | -0,717081171 | 0,221998475 | 3,230117561 |

| | | | | |
|---|---|---|---|---|
| ZC168.4 | 67,219783 | -0,7161476 | 0,2427856 | 2,949712 |
| R04A9.2 | 49,58857295 | -0,71549348 | 0,20641934 | 3,466213395 |
| C34F11.3 | 51,66880354 | -0,713676168 | 0,211564478 | 3,373327005 |
| D2023.2 | 110,5572416 | -0,704307925 | 0,169780862 | 4,148335198 |
| F20D12.5 | 147,844297 | -0,6614121 | 0,208454 | 3,172941 |
| C27C12.2 | 53,358324 | -0,6432961 | 0,2050648 | 3,137039 |
| B0379.4 | 113,4438214 | -0,641274152 | 0,194995714 | 3,288657677 |
| T22F3.3 | 95,10725985 | -0,636196605 | 0,170643472 | 3,728221161 |
| F33G12.6 | 49,705523 | -0,6355775 | 0,2051498 | 3,098114 |
| Y67D8C.10 | 133,6451705 | -0,580235468 | 0,164601144 | 3,525099839 |
| Y104H12BR.1 | 118,3102703 | -0,578900773 | 0,1741458 | 3,324230455 |
| F32D1.10 | 567,1404679 | -0,547717408 | 0,166800718 | 3,283663361 |
| C09B8.6 | 111,1363549 | -0,53931703 | 0,150108443 | 3,592849406 |
| F09C8.2 | 128,7068891 | -0,50641546 | 0,158541733 | 3,194209178 |
| H06O01.1 | 353,54509 | 0,3345586 | 0,1152489 | -2,902923 |
| E04A4.5 | 332,97673 | 0,3721427 | 0,1283115 | -2,900306 |
| Y54E2A.11 | 261,1276566 | 0,401590946 | 0,106728779 | -3,762724076 |
| C30C11.4 | 612,850295 | 0,4374191 | 0,1410602 | -3,100938 |
| C15H9.6 | 522,148047 | 0,4648205 | 0,1520833 | -3,056354 |
| F15C11.2 | 123,306095 | 0,526789417 | 0,162899086 | -3,233838994 |
| F53E10.6 | 64,851635 | 0,5352744 | 0,1784134 | -3,000193 |
| C08F8.1 | 57,347649 | 0,5571294 | 0,1915618 | -2,908354 |
| Y57G11C.15 | 615,380944 | 0,557488325 | 0,174102001 | -3,202078794 |
| F32D8.6 | 155,9616713 | 0,596410362 | 0,176885628 | -3,371728769 |
| F22B5.10 | 45,47339 | 0,6026689 | 0,200371 | -3,007766 |
| F15D4.3 | 138,472852 | 0,6049805 | 0,1974035 | -3,06469 |
| Y82E9BR.3 | 1184,618474 | 0,620688104 | 0,164772745 | -3,766934296 |
| T05E11.5 | 137,6151947 | 0,629322786 | 0,137295528 | -4,583709297 |
| F46H5.2 | 92,783169 | 0,6294835 | 0,2114079 | -2,977578 |
| M117.2 | 1072,493203 | 0,644787347 | 0,168339442 | -3,830280894 |
| Y38C1AA.4 | 86,1533312 | 0,666176373 | 0,180608895 | -3,688502568 |
| F25H2.10 | 787,551724 | 0,6854658 | 0,2339459 | -2,930019 |
| F01F1.8 | 1122,238472 | 0,6880134 | 0,2236028 | -3,076945 |
| Y22D7AL.5 | 1003,664268 | 0,7059176 | 0,2368409 | -2,980556 |
| T18H9.7 | 35,14044 | 0,7167174 | 0,2280709 | -3,14252 |
| Y48B6A.14 | 891,905536 | 0,7178651 | 0,2291007 | -3,133404 |
| T04G9.5 | 214,1718984 | 0,723360384 | 0,13470518 | -5,369952243 |
| C36E8.3 | 40,94627014 | 0,724220259 | 0,22443818 | -3,226813997 |
| Y54E10A.16 | 107,0887773 | 0,731731433 | 0,18997868 | -3,851650267 |
| F37C12.13 | 72,11643428 | 0,732758448 | 0,219450706 | -3,339057147 |
| C29F9.7 | 75,0175792 | 0,753545035 | 0,191438939 | -3,936216107 |
| ZC190.4 | 216,9294651 | 0,784720613 | 0,183963036 | -4,26564288 |
| F29C12.3 | 30,31078754 | 0,801940629 | 0,246449148 | -3,25398013 |
| F47B7.1 | 1051,288947 | 0,805641108 | 0,177838196 | -4,530191644 |
| R53.7 | 39,8465429 | 0,809794059 | 0,246127459 | -3,290141055 |
| C01G6.1 | 1019,26807 | 0,811666236 | 0,200492702 | -4,048358002 |
| T24H7.1 | 891,665078 | 0,8121701 | 0,2660965 | -3,052164 |
| T08B2.10 | 885,248143 | 0,8195439 | 0,2710568 | -3,023513 |
| F31E3.2 | 157,1369698 | 0,836505022 | 0,235089708 | -3,558237534 |
| K08C7.3 | 213,0354094 | 0,849643936 | 0,215937388 | -3,934677284 |

| | | | | |
|---|---|---|---|---|
| B0035.4 | 32,116959 | 0,8677055 | 0,2750063 | -3,155221 |
| T03F7.1 | 48,20000725 | 0,872908111 | 0,222642914 | -3,920664242 |
| C54E4.2 | 66,31447924 | 0,891449729 | 0,245486201 | -3,6313639 |
| C50F4.13 | 345,092877 | 0,9065933 | 0,3066341 | -2,956597 |
| T07H6.2 | 22,20730952 | 0,959874765 | 0,29685099 | -3,233523878 |
| T01C3.6 | 1621,348189 | 1,0369024 | 0,3554531 | -2,917129 |
| Y38A10A.5 | 339,1585075 | 1,040436754 | 0,26551121 | -3,918617046 |
| R13F6.4 | 127,4428287 | 1,064261695 | 0,235957075 | -4,51040382 |
| D1054.8 | 32,4693758 | 1,141138373 | 0,262148882 | -4,353016358 |
| T01A4.3 | 27,505256 | 1,1620759 | 0,3874103 | -2,9996 |
| F22F4.2 | 52,19754979 | 1,166371948 | 0,216036628 | -5,398954617 |
| T01G9.3 | 22,64063248 | 1,189535608 | 0,342317766 | -3,474945582 |
| T28F4.1 | 144,2926721 | 1,303271096 | 0,166783374 | -7,814154768 |
| C09D8.1 | 153,0201186 | 1,664274053 | 0,182705801 | -9,109037812 |
| F38B6.6 | 2,789607 | 2,8937525 | 0,9578417 | -3,021118 |
| F59B2.6 | 67,02389311 | 3,868209961 | 0,409688078 | -9,44184165 |

| pvalue | padj | Gene name |
|---|---|---|
| 3,03E-06 | 0,000520677 | F59D12.2 |
| 0,000143269 | 0,009029896 | arrd-26 |
| 7,75E-07 | 0,000152182 | glb-9 |
| 0,000452082 | 0,020985181 | F32D8.10 |
| 5,82E-06 | 0,000861384 | F43C9.2 |
| 0,000411553 | 0,019911065 | F10E9.12 |
| 1,48E-31 | 9,70E-28 | flp-15 |
| 2,25E-12 | 1,41E-09 | F31D4.8 |
| 8,76E-06 | 0,001094793 | cka-2 |
| 0,000615193 | 0,026580992 | T25F10.3 |
| 0,000398646 | 0,019562125 | C50D2.2 |
| 0,002819799 | 0,08043099 | nhr-14 |
| 1,12E-10 | 4,80E-08 | cdc-25.2 |
| 2,93E-06 | 0,000515652 | F35H10.10 |
| 0,000979698 | 0,03752634 | cah-6 |
| 0,000439583 | 0,020827138 | T01D3.7 |
| 0,000806283 | 0,033570674 | F55C12.4 |
| 0,001723041 | 0,05679789 | lron-9 |
| 0,003110319 | 0,08651603 | C49F8.3 |
| 1,18E-25 | 2,70E-22 | hil-2 |
| 1,56E-06 | 0,000296981 | klu-1 |
| 0,003825064 | 0,09850844 | bcmo-2 |
| 0,000355004 | 0,017932944 | grdn-1 |
| 0,000109105 | 0,007421331 | rege-1 |
| 0,002990216 | 0,0838481 | mua-6 |
| 0,000132181 | 0,008515241 | glb-33 |
| 0,00022471 | 0,012397753 | T05A8.3 |
| 0,001357223 | 0,047340432 | ptr-5 |
| 2,82E-31 | 9,70E-28 | R05H11.2 |
| 2,26E-10 | 9,15E-08 | glb-3 |
| 1,93E-05 | 0,001926314 | osta-2 |
| 2,37E-09 | 7,08E-07 | T09B4.5 |
| 3,42E-09 | 9,79E-07 | C05G5.7 |
| 0,000930504 | 0,036321382 | srap-1 |
| 0,000231257 | 0,012609035 | T04F8.9 |
| 1,47E-05 | 0,001625322 | F15G9.5 |
| 0,000815112 | 0,033605025 | C30G7.2 |
| 1,87E-05 | 0,001919869 | hum-5 |
| 0,00014609 | 0,009123996 | sma-10 |
| 8,28E-05 | 0,006154084 | T05A8.8 |
| 3,22E-05 | 0,002912655 | fasn-1 |
| 0,002126745 | 0,06581414 | rimb-1 |
| 8,54E-08 | 1,89E-05 | pqn-48 |
| 0,003497502 | 0,09399201 | T19D12.6 |
| 0,003165864 | 0,08740584 | sms-3 |
| 1,32E-11 | 7,33E-09 | F19B10.10 |
| 0,002328636 | 0,06955533 | ces-2 |
| 0,002560174 | 0,07484424 | C54G4.5 |

| | | |
|---|---|---|
| 0,002186709 | 0,06706558 | ctbp-1 |
| 0,000994606 | 0,03752634 | C32D5.7 |
| 1,60E-11 | 7,85E-09 | lec-4 |
| 9,97E-09 | 2,74E-06 | drag-1 |
| 0,000337958 | 0,017346596 | scrm-3 |
| 0,001064448 | 0,038897648 | F45E1.2 |
| 0,000116103 | 0,007819868 | D2092.6 |
| 0,001799672 | 0,05859595 | nhr-25 |
| 0,000965333 | 0,037257512 | che-7 |
| 0,000821782 | 0,033605025 | W08A12.3 |
| 0,000383677 | 0,01910045 | lim-9 |
| 9,75E-21 | 1,12E-17 | mls-2 |
| 1,60E-05 | 0,001686888 | let-522 |
| 4,46E-18 | 3,83E-15 | ras-1 |
| 0,000191317 | 0,011039868 | R08C7.12 |
| 0,000196299 | 0,011063091 | plc-3 |
| 0,003599772 | 0,0958544 | twk-24 |
| 0,002788729 | 0,08016136 | T19E7.6 |
| 0,00083819 | 0,03397145 | F08G2.7 |
| 0,000338347 | 0,017346596 | R02E12.5 |
| 3,18E-05 | 0,002912655 | ZC132.4 |
| 1,39E-11 | 7,33E-09 | efl-3 |
| 0,000545517 | 0,024178722 | nhr-97 |
| 4,23E-05 | 0,003635702 | Y65B4BL.1 |
| 5,50E-05 | 0,004498063 | glb-16 |
| 0,001482056 | 0,05055851 | snn-1 |
| 0,000449663 | 0,020985181 | unc-36 |
| 0,002051405 | 0,06405978 | pgp-2 |
| 0,003175232 | 0,08740584 | F19B2.5 |
| 1,90E-05 | 0,001919869 | rgl-1 |
| 4,18E-05 | 0,003633237 | fhod-1 |
| 1,89E-06 | 0,000350778 | vab-8 |
| 0,000502476 | 0,02316784 | vab-9 |
| 5,16E-05 | 0,004272492 | nlf-1 |
| 0,003828494 | 0,09850844 | T13H5.6 |
| 0,001856905 | 0,06017422 | dep-1 |
| 3,69E-05 | 0,003248792 | F21D5.9 |
| 0,002521144 | 0,07401821 | F58H10.1 |
| 7,10E-10 | 2,44E-07 | hil-3 |
| 0,001007888 | 0,03752634 | F44E2.10 |
| 0,000300262 | 0,015746539 | nac-3 |
| 7,14E-08 | 1,69E-05 | unc-98 |
| 5,29E-16 | 4,04E-13 | T10B10.4 |
| 0,001486582 | 0,05055851 | C13C4.4 |
| 4,58E-10 | 1,66E-07 | F07C6.4 |
| 7,35E-05 | 0,005670393 | zyx-1 |
| 2,82E-05 | 0,002619789 | F37A8.5 |
| 1,52E-24 | 2,61E-21 | W02D9.10 |
| 0,000142513 | 0,009029896 | W02B8.1 |
| 9,02E-05 | 0,006387679 | tsp-8 |

| | | |
|---|---|---|
| 0,000171106 | 0,010221725 | unc-9 |
| 2,03E-05 | 0,001960523 | rhi-1 |
| 0,000572296 | 0,025203036 | let-2 |
| 0,00090668 | 0,035798239 | cah-3 |
| 0,000188988 | 0,011002922 | sdpn-1 |
| 7,14E-06 | 0,000962185 | smp-1 |
| 0,000406788 | 0,019820111 | egl-19 |
| 0,001462427 | 0,05023438 | F26F4.5 |
| 0,000179004 | 0,010601354 | shn-1 |
| 6,39E-05 | 0,00504642 | T12A7.2 |
| 0,001278597 | 0,04527815 | jph-1 |
| 2,04E-08 | 5,40E-06 | Y43F8B.2 |
| 2,73E-05 | 0,002565556 | flp-10 |
| 8,63E-06 | 0,001094793 | gon-1 |
| 0,00230605 | 0,06950667 | sph-1 |
| 5,12E-06 | 0,000781728 | Y15E3A.5 |
| 0,000927719 | 0,036321382 | dhs-19 |
| 1,31E-05 | 0,001552048 | mex-3 |
| 2,22E-06 | 0,000401889 | flh-2 |
| 0,000129075 | 0,00844519 | nrfl-1 |
| 2,34E-08 | 5,95E-06 | rpy-1 |
| 7,90E-06 | 0,001044211 | Y71H10B.1 |
| 6,32E-05 | 0,00504638 | tlp-1 |
| 1,19E-05 | 0,001438074 | acl-12 |
| 0,001966818 | 0,06253367 | casy-1 |
| 0,000106002 | 0,007355931 | ina-1 |
| 3,22E-06 | 0,000539572 | hxk-1 |
| 8,07E-06 | 0,001046228 | bre-3 |
| 5,44E-07 | 0,000109845 | C02B10.5 |
| 2,18E-07 | 4,55E-05 | F52D2.6 |
| 3,73E-06 | 0,000595983 | ric-4 |
| 0,003657358 | 0,09663866 | T19H5.4 |
| 1,87E-11 | 8,57E-09 | gpa-4 |
| 0,002316889 | 0,06950667 | unc-22 |
| 0,003294336 | 0,09016769 | hsp-70 |
| 2,88E-10 | 1,10E-07 | snf-3 |
| 0,000446881 | 0,020985181 | obr-1 |
| 0,000164015 | 0,010047495 | snf-1 |
| 0,001977672 | 0,06253367 | mop-25.1 |
| 0,000629373 | 0,027023689 | H24G06.1 |
| 1,79E-05 | 0,001863254 | nfki-1 |
| 7,09E-05 | 0,005536554 | T05H10.4 |
| 1,97E-09 | 6,14E-07 | sup-1 |
| 0,000260651 | 0,014099764 | C17C3.1 |
| 0,002310384 | 0,06950667 | cyb-3 |
| 0,000531714 | 0,023719975 | svop-1 |
| 1,04E-05 | 0,001275857 | abts-4 |
| 1,58E-05 | 0,001686888 | aspm-1 |
| 0,000882972 | 0,035208907 | unc-13 |
| 1,85E-09 | 6,04E-07 | W09D6.5 |

| | | |
|---|---|---|
| 8,74E-05 | 0,006373247 | pkc-1 |
| 0,00096106 | 0,037257512 | eat-6 |
| 9,42E-05 | 0,006605343 | peb-1 |
| 1,40E-05 | 0,001573732 | K08H10.2 |
| 0,00015105 | 0,009348777 | klp-18 |
| 0,000821315 | 0,033605025 | tra-1 |
| 0,003683266 | 0,0969397 | C53C11.5 |
| 0,000531371 | 0,023719975 | ung-1 |
| 0,001364261 | 0,047340432 | lin-42 |
| 0,00343769 | 0,09334756 | scpl-3 |
| 0,003487906 | 0,09399201 | sipa-1 |
| 0,002429234 | 0,07224606 | C07A9.12 |
| 2,67E-05 | 0,002543651 | arrd-22 |
| 6,02E-06 | 0,000861598 | lit-1 |
| 0,000391153 | 0,019332537 | soc-2 |
| 1,48E-07 | 3,17E-05 | F55F1.1 |
| 0,00378633 | 0,09815882 | jac-1 |
| 0,001749514 | 0,05723411 | Y53G8AM.8 |
| 0,000849093 | 0,034112697 | unc-87 |
| 4,61E-05 | 0,003910061 | C14B4.2 |
| 0,001966142 | 0,06253367 | ptl-1 |
| 0,0020434 | 0,06405978 | tth-1 |
| 3,50E-06 | 0,000572629 | prkl-1 |
| 0,003502468 | 0,09399201 | wrm-1 |
| 0,000984851 | 0,03752634 | nab-1 |
| 1,37E-05 | 0,001565421 | atn-1 |
| 0,000434763 | 0,020741815 | mct-6 |
| 0,000518232 | 0,023577821 | C35D10.13 |
| 0,000286546 | 0,015142857 | emb-9 |
| 0,000669858 | 0,028406926 | F52G3.1 |
| 6,53E-06 | 0,000896852 | fzy-1 |
| 0,000196462 | 0,011063091 | his-72 |
| 0,003752447 | 0,09764889 | vab-10 |
| 0,001543442 | 0,05197767 | swsn-1 |
| 0,002821524 | 0,08043099 | cpn-1 |
| 8,05E-05 | 0,006141463 | unc-84 |
| 0,000576193 | 0,025213016 | catp-7 |
| 0,00102733 | 0,037742019 | epg-9 |
| 0,000612968 | 0,026580992 | ima-2 |
| 0,000274271 | 0,014720628 | hum-2 |
| 0,000642134 | 0,027400384 | rod-1 |
| 0,001984329 | 0,06253367 | bmk-1 |
| 1,59E-05 | 0,001686888 | cdk-1 |
| 0,001010535 | 0,03752634 | ZK484.4 |
| 0,000132625 | 0,008515241 | arrd-25 |
| 0,000210409 | 0,011752131 | maph-1.1 |
| 0,000187794 | 0,011002922 | mdf-2 |
| 0,00034227 | 0,01741773 | dcp-66 |
| 0,001285621 | 0,045293417 | F28C1.3 |
| 0,001237393 | 0,044275478 | T09B9.4 |

| | | |
|---|---|---|
| 0,003180707 | 0,08740584 | cyb-1 |
| 0,000527844 | 0,023719975 | nrde-3 |
| 0,000742657 | 0,031291163 | ampd-1 |
| 3,35E-05 | 0,00298802 | pyc-1 |
| 0,001509032 | 0,05106922 | exc-9 |
| 0,001706635 | 0,05664048 | egrh-1 |
| 0,001006664 | 0,03752634 | scpl-1 |
| 0,000192836 | 0,011039868 | pygl-1 |
| 0,001947568 | 0,06252238 | F33G12.6 |
| 0,000423323 | 0,020337276 | mca-3 |
| 0,000886629 | 0,035208907 | plst-1 |
| 0,001024672 | 0,037742019 | mcm-7 |
| 0,000327082 | 0,017023112 | hsp-25 |
| 0,001402145 | 0,048405702 | F09C8.2 |
| 0,003696973 | 0,0969397 | pdi-3 |
| 0,00372798 | 0,09738108 | timm-17B.1 |
| 0,000168073 | 0,010128586 | eif-3.B |
| 0,001929085 | 0,06221977 | hsp-110 |
| 0,002240467 | 0,06840894 | hsp-3 |
| 0,001221383 | 0,043979911 | ubql-1 |
| 0,002698089 | 0,07803344 | F53E10.6 |
| 0,003633372 | 0,09637554 | pfd-1 |
| 0,001364397 | 0,047340432 | sec-61 |
| 0,00074698 | 0,031291163 | emo-1 |
| 0,00263176 | 0,07661097 | F22B5.10 |
| 0,002178956 | 0,06706558 | romo-1 |
| 0,000165264 | 0,010047495 | Y82E9BR.3 |
| 4,57E-06 | 0,00071323 | imp-2 |
| 0,002905354 | 0,08180238 | F46H5.2 |
| 0,000127997 | 0,00844519 | par-5 |
| 0,000225578 | 0,012397753 | tcl-2 |
| 0,003389412 | 0,09240184 | rla-0 |
| 0,002091341 | 0,06501137 | cct-6 |
| 0,002877257 | 0,08134467 | hsp-60 |
| 0,001675004 | 0,05586057 | tag-232 |
| 0,001727913 | 0,05679789 | hmg-1.1 |
| 7,88E-08 | 1,80E-05 | trap-2 |
| 0,001251768 | 0,044557753 | pxd-1 |
| 0,000117325 | 0,00782543 | mab-31 |
| 0,000840633 | 0,03397145 | exos-9 |
| 8,28E-05 | 0,006154084 | pat-4 |
| 1,99E-05 | 0,001956256 | ZC190.4 |
| 0,001138002 | 0,041365484 | rict-1 |
| 5,89E-06 | 0,000861384 | F47B7.1 |
| 0,001001372 | 0,03752634 | aakg-5 |
| 5,16E-05 | 0,004272492 | aqp-2 |
| 0,002271978 | 0,0690641 | phb-2 |
| 0,002498583 | 0,0739882 | rps-17 |
| 0,000373352 | 0,018722084 | F31E3.2 |
| 8,33E-05 | 0,006154084 | epi-1 |

| | | |
|---|---|---|
| 0,001603765 | 0,05374569 | pfd-4 |
| 8,83E-05 | 0,006373247 | snf-11 |
| 0,000281927 | 0,015014269 | test-1 |
| 0,003110547 | 0,08651603 | his-35 |
| 0,001222731 | 0,043979911 | mom-1 |
| 0,0035327 | 0,09443442 | rps-16 |
| 8,91E-05 | 0,006373247 | crt-1 |
| 6,47E-06 | 0,000896852 | ten-1 |
| 1,34E-05 | 0,001563532 | D1054.8 |
| 0,002703342 | 0,07803344 | T01A4.3 |
| 6,70E-08 | 1,64E-05 | inx-3 |
| 0,000510957 | 0,02340183 | dma-1 |
| 5,53E-15 | 3,80E-12 | T28F4.1 |
| 8,31E-20 | 8,16E-17 | ptp-3 |
| 0,002518435 | 0,07401821 | F38B6.6 |
| 3,66E-21 | 5,03E-18 | zif-1 |
